# Voxelwise encoding models show that cerebellar language representations are highly conceptual

**DOI:** 10.1101/2021.01.18.427158

**Authors:** Amanda LeBel, Shailee Jain, Alexander G. Huth

**Affiliations:** Helen Wills Neuroscience Institute, University of California Berkeley, Berkeley, CA 94720, USA; Department of Neuroscience, The University of Texas at Austin, Austin, TX 78712, USA; Department of Computer Science; The University of Texas at Austin, Austin, TX 78712, USA

## Abstract

There is a growing body of research demonstrating that the cerebellum is involved in language understanding. Early theories assumed that the cerebellum is involved in low-level language processing. However, those theories are at odds with recent work demonstrating cerebellar activation during cognitive tasks. Using natural language stimuli and an encoding model framework, we performed an fMRI experiment where subjects passively listened to five hours of natural language stimuli which allowed us to analyze language processing in the cerebellum with higher precision than previous work. We used this data to fit voxelwise encoding models with five different feature spaces that span the hierarchy of language processing from acoustic input to high-level conceptual processing. Examining the prediction performance of these models on separate BOLD data shows that cerebellar responses to language are almost entirely explained by high-level conceptual language features rather than low-level acoustic or phonemic features. Additionally, we found that the cerebellum has a higher proportion of voxels that represent social semantic categories, which include “social” and “people” words, and lower representations of all other semantic categories, including “mental”, “concrete”, and “place” words, than cortex. This suggests that the cerebellum is representing language at a conceptual level with a preference for social information.

**Significance Statement:** Recent work has demonstrated that, beyond its typical role in motor planning, the cerebellum is implicated in a wide variety of tasks including language. However, little is known about the language representations in the cerebellum, or how those representations compare to cortex. Using voxelwise encoding models and natural language fMRI data, we demonstrate here that language representations are significantly different in the cerebellum as compared to cortex. Cerebellum language representations are almost entirely semantic, and the cerebellum contains over-representation of social semantic information as compared to cortex. These results suggest that the cerebellum is not involved in language processing per se, but cognitive processing more generally.

## Introduction

The cerebellum is known to be involved in a diverse set of cognitive processes including attention(Allen et al., 1997), working memory(Brissenden et al., 2018), object recognition(Liu et al., 1999), and language processing(Booth et al., 2007; Stoodley & Schmahmann, 2009). Evidence for the cognitive function of the cerebellum in healthy subjects has come largely from neuroimaging studies, which have found that certain cognitive tasks elicit consistently localized BOLD responses across cerebellum(King et al., 2018) and that resting-state BOLD fluctuations in cerebellum align to known resting-state networks in cortex(Buckner et al., 2011; Marek et al., 2018). However, little is known about what role the cerebellum plays in cognitive processes, or how representations in the cerebellum might differ from those found in cortex.

Language understanding is a highly complex cognitive process, which makes it a rich area of research to study cognitive processing. Hierarchically organized networks for language processing are widely distributed across much of cortex(J. R. Binder et al., 1997; de Heer et al., 2017; Dronkers et al., 2004; Hickok & Poeppel, 2007; Poeppel et al., 2012). These networks include some putative “language specific” areas in temporal and inferior frontal cortex(Fedorenko et al., 2011), as well as non-language specific conceptual areas in temporal, parietal, and prefrontal cortex(Fedorenko et al., 2013). However, it is unclear whether these networks are also reflected in the cerebellum. Clinical evidence for a cerebellar role in language processing is found in work on cerebellar cognitive affective syndrome (CCAS), which shows that patients with acquired cerebellar damage experience language degradation which can include agrammatism, dysprosody, and anomia(Schmahmann & Sherman, 1998). However, the subtlety and variability of these effects have made it difficult to form a complete picture. Early work into language deficits from cerebellar lesions has often conflicted with cases suggesting a degradation in grammar while preserving semantic content(Frank et al., 2008; Justus, 2004; M. C. Silveri et al., 1994) and other work suggesting a more uniform degradation in language processing that includes semantic content(Cook et al., 2004; Fiez et al., 1992; Maria Caterina Silveri & Misciagna, 2000). However, it is unclear if the standard aphasia tests used in these studies are sensitive enough to detect deficits from cerebellar damage (Cook et al., 2004; Murdoch, 2010). Our goal is to determine how language perception is localized in the cerebellum, what aspects of language are represented in the cerebellum, and how this compares to language processing systems in cortex.

Here we modeled cortical and cerebellar representations of natural speech using three different categories of features that span the putative language processing hierarchy (de Heer et al., 2017; Hickok & Poeppel, 2007): modality-specific, language-specific, and conceptual. Modality-specific features capture information specific to how people perceive the language stimulus. In this study, subjects listened to audio recordings of naturally spoken narrative stories, so we used a feature space that captures frequency information in sound(Cheung et al., 2016). This feature space is known to be represented in auditory cortex(de Heer et al., 2017). Building upon modality-specific features, language-specific features capture information that only exists in language, like phoneme articulations and syntax. These feature spaces are known to be represented in STG(de Heer et al., 2017; Fedorenko et al., 2011) and inferior frontal cortex(de Heer et al., 2017). Finally, conceptual features capture information about the meaning conveyed by language, which is known to be represented across broad regions of cortex, overlapping with other cognitive tasks(de Heer et al., 2017; Fedorenko et al., 2013). Previous work used similar methods to demonstrate that there is a hierarchy across these feature categories in cortex, where modality-specific information feeds into language-specific and then conceptual representations(de Heer et al., 2017). Here we investigated whether this hierarchy is replicated in the cerebellum, or if the cerebellum is specifically involved in only some aspects of language processing. For ease of language, “cortex” here refers exclusively to the cerebral cortex and “cerebellum” refers to the whole cerebellum, as cerebellar white matter was not excluded from analysis.

To determine which aspects of language the cerebellum is involved in processing or representing, we conducted a functional MRI (fMRI) experiment where subjects passively listened to 27 natural, narrative stories (5.4 hours) about a diverse set of topics. We then used voxelwise encoding models (**Figure 1**) to determine how well each set of speech-related features could predict each voxel in each subject. The stimuli were first transformed into 5 different feature spaces: spectral, articulatory, part-of-speech, word-level semantic, and context-level (multi-word) semantic. We used ridge regression to fit voxelwise encoding models with each feature space, and then tested how well these encoding models could predict responses to a new story that was not used for model fitting. Finally, we used variance partitioning to measure how much variance in cerebellar and cortical BOLD responses is uniquely explained by each of the five feature spaces. We found substantial evidence that the cerebellum represents language at a high conceptual and semantic level, and no strong evidence that the cerebellum represents any language-specific or modality-specific information.

**Figure 1.**
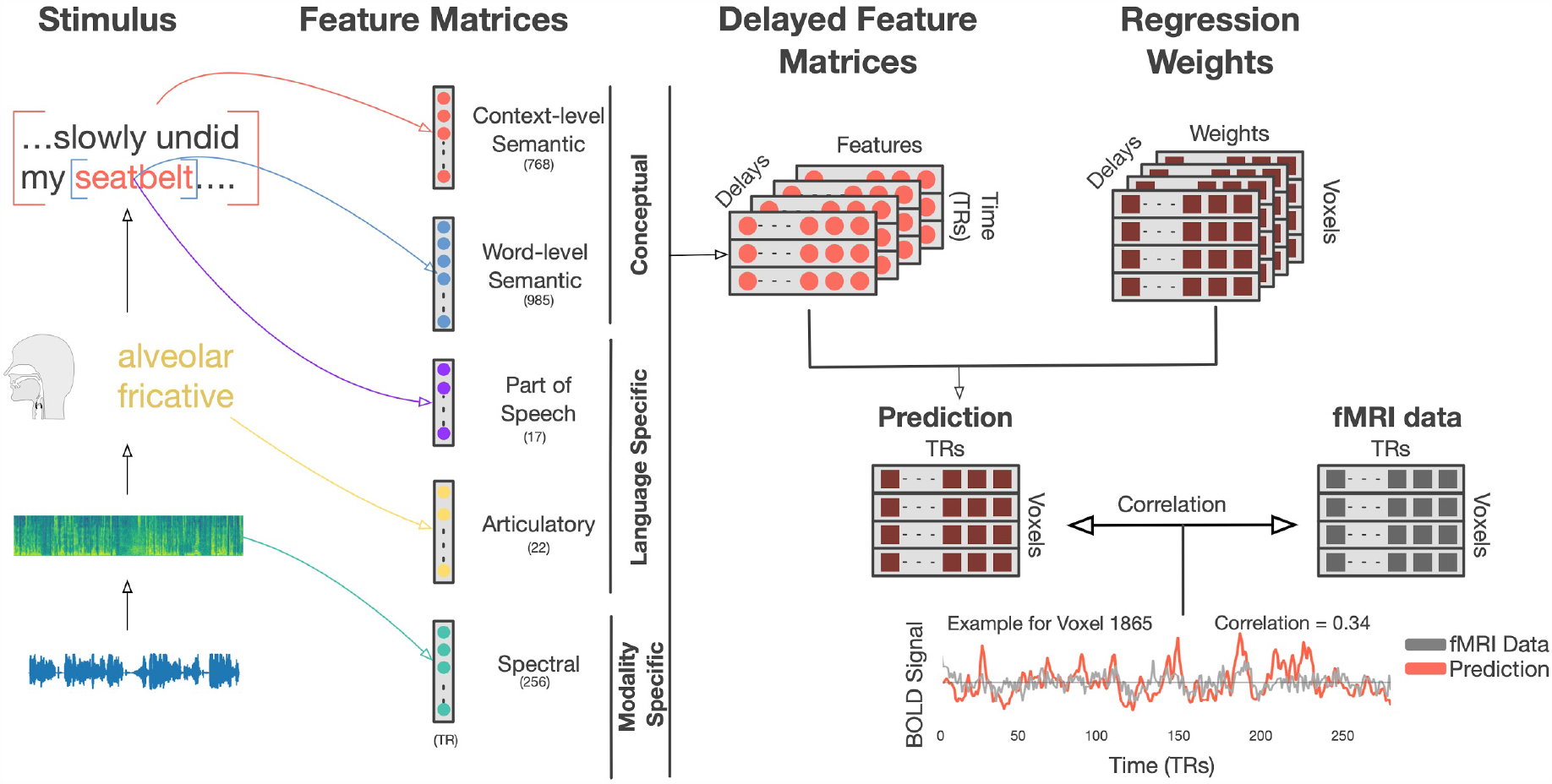
Voxelwise encoding model construction. To localize different stages of language processing across the cerebellum we used five feature spaces to predict voxelwise BOLD responses in each subject: spectral, articulatory, part-of-speech, word-level semantic, and context-level semantic. Each 10-15 minute stimulus story was transcribed and temporally aligned to the audio recording at the word and phoneme level. Features were then extracted for each of the five feature spaces. The features for the spectral model are 256 bands of a mel-frequency spectrogram, the features for the articulatory model are a 22 length n-hot vector, the features for the part-of-speech model are a 1-hot 17 length vector, the features for the word-level semantic model are a 985-dimensional vector based on statistical word co-occurrence, and the features for the context-level semantic model are a 768 dimensional vector based on GPT(Radford et al., 2018), a neural network language model that incorporates context (preceding words) into the representation of the current word. Features were extracted for each timepoint, word, or phoneme, and concatenated into a feature matrix. The feature matrix was then resampled to the rate of the BOLD signal (0.5 Hz) and delayed to form an FIR model that accounts for hemodynamics. Then regularized linear regression was used to fit weights that predict each voxel’s BOLD signal from the stimulus matrix. Finally, models were used to predict responses on a held out test dataset that was not used for model fitting. Model performance was assessed as the linear correlation between held out BOLD data and model predictions for each voxel.

In addition, we used the word-level semantic encoding models to determine whether the cerebellum represents different semantic categories than cortex. This analysis showed that all semantic categories are represented in both the cerebellum and cortex, but that the cerebellum has an overrepresentation of social semantic categories and an underrepresentation of mental, concrete, and place-related semantic categories as compared to cortex.

## Methods

### Participants

Data was collected from three male subjects and two female subjects: UT-S-01(female, age=24), UT-S-02 (male, age 34), UT-S-06 (female, age 23), UT-S-07 (male, age=25), UT-S-08 (male, age=24). Three of the subjects were authors (UT-S-01:S.J, UT-S-02:A.G.H, and UT-S-06:A.L). All subjects were healthy and had normal hearing. The experimental protocol was approved by the Institutional Review Board at the University of Texas at Austin. Written informed consent was obtained from all subjects.

### fMRI collection

MRI data was collected on a 3T Siemens Skyra scanner at the UT Austin Biomedical Imaging Center using a 64 channel Siemens volume coil. Functional scans were collected using gradient echo EPI with repetition time (TR) = 2.00 s, echo time (TE) = 30.8 ms, flip angle = 71°, multi-band factor (simultaneous multi-slice) = 2, voxel size = 2.6mm x 2.6mm x 2.6mm (slice thickness = 2.6mm), matrix size = (84, 84), and field of view = 220 mm. Field of view covered both the cortex and the cerebellum in their entirety for all subjects. Anatomical scans were collected using a T1-weighted multi-echo MP-RAGE sequence on the same 3T scanner with voxel size = 1mm x 1mm x 1mm following the Freesurfer morphometry protocol. Anatomical data for subject UT-S-02 was collected on a 3T Siemens TIM Trio at the Berkeley Brain Imaging Center with a 32-Channel Seimen’s volume coil using the same sequence.

Known regions of interest (ROIs) were localized separately in each subject. Three different tasks were used to define ROIs, these include a visual category localizer, an auditory cortex localizer, and a motor localizer.

For the visual category localizer, data were collected in six 4.5 minute scans consisting of 16 blocks of 16 seconds each. During each block 20 images of either places, faces, bodies, household objects, or spatially scrambled objects were displayed. Subjects were asked to pay attention for the same image being presented twice in a row. The corresponding ROIs defined in cortex with this localizer were the fusiform face area (FFA)(Kanwisher et al., 1997), occipital face area (OFA)(Kanwisher et al., 1997), extrastriate body area (EBA)(Downing et al., 2001), parahippocampal place area (PPA)(Epstein & Kanwisher, 1998), and the occipital place area (OPA).

Motor localizer data were collected during 2 identical 10-minute scans. The subject was cued to perform six different tasks in a random order in 20-second blocks. The cues were ‘hand’, ‘foot’, ‘mouth’, ‘speak’, saccade, and ‘rest’ presented as a word at the center of the screen, except for the saccade cue which was presented as a random array of dots. For the hand cue subjects were instructed to make small finger-drumming movements for the entirety of the time the cue was displayed. For the foot cue, the subjects were instructed to make small foot and toe movements. For the mouth cue, subjects were instructed to make small vocalizations that were nonsense syllables such as *balabalabala*. For the speak cue, subjects were instructed to self-generate a narrative without vocalization. For the saccade cue, subjects were instructed to look around for the duration of the task.

Weight maps for the motor areas were used to define primary motor and somatosensory areas for the hands, feet, and mouth; supplemental motor areas for the hands and feet, secondary motor areas for the hands, feet, and mouth, the ventral premotor hand area. The weight map for the saccade responses was used to define the frontal eye field and intraparietal sulcus visual areas. The weight map for the speech production was used to define broca’s area and the superior ventral premotor area (sPMV) speech area(Chang et al., 2011). In the cerebellum, weight maps for each subject were resliced in SUIT space(Diedrichsen, 2006) and then the resliced maps were averaged across subjects for each task. Motor areas for the hand, mouth, foot, and saccade tasks were defined in the posterior and anterior lobe.

Auditory cortex localizer data were collected in one 10-minute scan. The subject listened to 10 repeats of 1-minute auditory stimulus each containing 20 seconds of music (Arcade Fire), speech (Ira Glass, This American Life), and natural sound (a babbling brook). To determine whether a voxel was responsive to auditory stimulus, the repeatability of the voxel response across the 10 repeats was calculated using an *F-*statistic. This map was used to define the auditory cortex (AC).

### fMRI preprocessing

All functional data were motion corrected using the FMRIB Linear Image Registration Tool (FLIRT) from FSL 5.0(Woolrich et al., 2009). FLIRT was used to align all data to a template that was made from the average of all functional runs in the first story session for each subject. These automatic alignments were manually checked. Low frequency voxel response drift was identified using a 2^nd^ order Savitzky-Golay filter with a 120 second window and then subtracted from the signal. To avoid artifacts from onsets and poor detrending performance, responses were trimmed by removing 20 seconds (10 volumes) at the beginning and end of each scan. This removed the 10-second silent period as well as the first and last 10 seconds of each story. The mean response for each voxel was subtracted and the remaining response was scaled to have unit variance.

### Cortical Surface reconstruction and Visualization

For cortical surfaces, meshes were generated from the T1-weighted anatomical scans using freesurfer(Dale et al., 1999). Before surface reconstruction, anatomical surface segmentations were hand-checked and corrected. Blender was used to remove the corpus callosum and make relaxation cuts for flattening. Functional images were aligned to the cortical surface using boundary based registration (BBR) implemented in FSL. These were checked for accuracy and adjustments were made as necessary.

For the cerebellum cortical surfaces, the SUIT toolbox(Diedrichsen, 2006) was used to isolate the cerebellum from the rest of the brain using the T1-weighted anatomical image. The anatomical maps for the cerebellum were normalized into SUIT space using the SUIT registration algorithm. After encoding model fitting, cerebellar functional results were transformed into anatomical space and then resliced using SUIT. The SUIT flatmap and surface was added to the pycortex database for the purpose of surface visualization.

Model maps were created by projecting the values for each voxel onto the cortical surface using the ‘nearest’ scheme in pycortex software(Gao et al., 2015). This projection finds the location of each pixel in the image in 3D space, and assigns that pixel the associated value.

### Stimulus set

The modeling training stimulus set consisted of 26 10-15 min stories taken from *The Moth Radio Hour*. In each story, a single speaker tells an autobiographical story without reading from a prepared speech. Each story was played during one scan with a buffer of 10 seconds on either side of the story start and stop. Data collection was broken up into 6 different days, the first session involving the anatomical scan and localizers, and each successive session consisting of 4 to 5 stories, plus one additional story used for model prediction. This additional story (which was not one of the 26 stories used for model training) was played in every session and the responses to this story were averaged. Stories were played over Sensimetrics S14 in-ear piezoelectric headphones. The audio for each story was filtered to correct for frequency response and phase errors induced by the headphones using calibration data provided by sensimetrics and custom python code (https://github.com/alexhuth/sensimetrics_filter). All stimuli were played at 44.1 kHz using the pygame library in Python.

Each story was manually transcribed by one listener. Certain sounds (for example, laughter and breathing) were also marked to improve the accuracy of the automated alignment. The audio of each story was downsampled to 11kHz and the Penn Phonetics Lab Forced Aligner (P2FA)(Jiahong Yuan, 2008) was used to automatically align the audio to the transcript. Praat(P Boersma, 2014) was then used to check and correct each aligned transcript manually.

### Feature Spaces

Five feature spaces were used to cover the hierarchy of language processing. Each feature space was fit separately for each subject. The spectral feature space was a mel-band spectrogram(Jiahong Yuan, 2008) with frequencies ranging from approximately 0 Hz to 8kHz with 256 windows. The articulatory feature space was a n-hot feature space where each phoneme is assigned a 1 for each articulation that is required to produce the sound and a 0 for every other articulation for a total of 22 features per phoneme. For the part-of-speech feature space, a one-hot vector of 17 features was assigned to each word noting the part of speech for each word in each story. Part of speech tagging was done using the flair package(Akbik et al., 2019). Flair is a language model that uses recurrent neural networks to tag speech into 17 categories (e.g. noun, verb, number, determiner, etc). The word-level semantic space was a 985-dimensional feature space based on word co-occurrence(Huth et al., 2016). Each word in the stimulus set was assigned the vector associated with it in the original space. If the word in the story was not present in the original semantic space, it was assigned a vector of length 985 of zeros. The contextual semantic space was based on a fine-tuned GPT language model(Radford et al., n.d.). GPT is a state of the art language model that takes into account previous words while generating features for the current word. To assign features to each word, we extracted 768-dimensional feature vectors from layer 9 with a context length of 25 words. We chose layer 9 because it is a midlayer of GPT and it has been demonstrated that middle layers of recurrent language models are best able to predict brain activity(Jain & Huth, 2018; Toneva & Wehbe, 2019).

### Experimental Design and Statistical Analysis

#### Encoding model fitting

We used each of the 5 features spaces to fit a linearized finite impulse response (FIR) model to every cortical voxel in each subject. The cerebellar models and the cortical models were fit separately. The stimulus matrix for each story was downsampled using a 3-lobe Lanczos filter, then z-scored and concatenated together. To fit the linear model the stimulus matrix has to account for variance in the hemodynamic response function across voxels. To do this we concatenate 4 delayed copies of the stimulus (using delays of 1, 2, 3, and 4 time points). This final stimulus matrix is then regressed with the BOLD data using ridge regression. We then test the model using a held out data set. This is done by taking the dot product of the weight matrix from the regression with the stimulus matrix from the held out test set, resulting in a voxel by timepoint matrix. This resulting matrix is compared to the actual BOLD data for the held out test set and the correlation calculated over time for each voxel to give a measure of model performance. The correlation was then noise-ceiling corrected for some analyses (noted in the text)(Schoppe et al., 2016). Total model performance metrics were computed using the mean R^2^ across voxels. Mean was used instead of summation to better account for the difference in number of voxels over the cerebellum as compared to the cortex. To keep the scale of the weights consistent, a single value of the regularization coefficient was used for all voxels in both the cerebellum and cortex in all subjects. To find the best regularization coefficient, the regression procedure was bootstrapped 50 times in each subject and a regularization performance curve was obtained for each subject by averaging the bootstrap sample correlations across the 50 samples, then across voxels, and finally across the 6 subjects and the best overall value of the regularization parameter was selected. This was done separately for each feature space.

### Individual model comparison

Encoding models with each of the feature spaces were fit in the cerebellum and cortex in each subject and the regression weights were used to predict a held out test set. Model performance for each voxel was estimated by taking the correlation of the predicted time series for each voxel with the actual data. Then to test if the model performance was significant the time series for each voxel was randomly shuffled in blocks of TRs and the correlation with the predicted time series was recalculated. This was done for 10000 permutations to gain a null distribution of responses. Lastly, the Fisher-corrected p-value was calculated and this was FDR corrected to account for all the comparisons. A threshold of *q*(*FDR*) < 0.05 was used to test for significantly well predicted voxels. This was done individually in the cerebellum and cortex in each subject for each model. The correlations were also noise-ceiling corrected. Comparison was done across subjects by taking the average R^2^ of all voxels in each subject in the cerebellum and cortex

### Variance Partitioning

Because 5-way variance partitioning has too many partitions to be interpretable, we used two versions of variance partitioning to test specific hypotheses. The first version looked at the unique variance explained by each model. This was done to test if one feature space is uniquely better at predicting cerebellar or cortical voxels. The second version was a pairwise variance partitioning where each model was jointly fit with the contextual semantic space. This was done to test for the specific hypothesis that the contextual semantic model is better predicting the same areas as the low level models in the cerebellum, i.e. are there unique low-level language representations in the cerebellum or is the contextual semantic model better predicting the same areas as the low-level models. To do variance partitioning, joint models with the concatenated feature spaces are fit and then used to predict the held out data set. To be succinct, the variance explained by the five feature spaces will be written as sets A-E.

Unique Partition - The following nested models were fit as follows

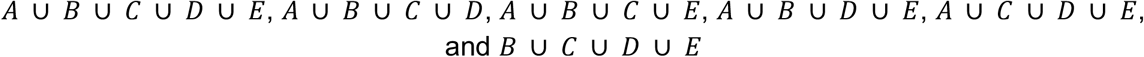

The variance uniquely explained by each feature space without any overlap from the other feature spaces, or relative complement (RC), was then calculated for each feature space as follows:

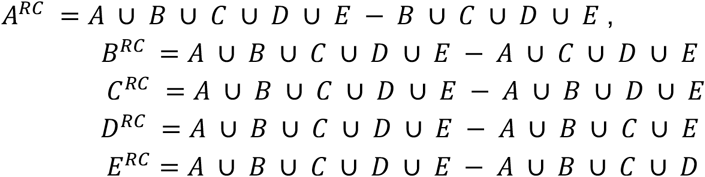

A Fisher-corrected permutation test with 10000 permutations was done in each subject in both the cerebellum and cortex for each voxel for the unique partitions using the joint *A*⋃*B*⋃*C* ⋃ *D*⋃*E* model. Multiple comparison correction was done using FDR with a threshold of *p* < 0.0*5*. Cerebellar data was resliced after the calculation of the unique partitions and the significance testing. The mean of the variance explained was calculated for each subject, in each partition, in the cerebellum and cortex.

Pairwise Variance Partitioning - The following concatenated models were fit as follows (where A is the contextual semantic feature space):

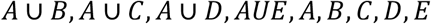

The variance explained by the intersections were calculated as follows:

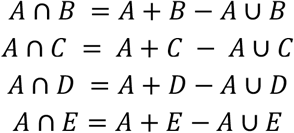

Then the unique contribution of each feature space in each pair can be calculated. This is the unique contribution without overlap from the other feature space noted as RC/X where X is the other paired feature space. These are calculated as follows:

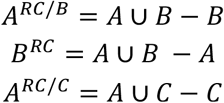

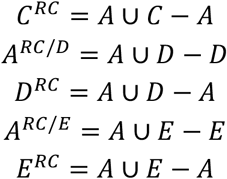

A Fisher-corrected permutation test with 10000 permutations was done in each subject in both the cerebellum and cortex for each voxel for the unique partitions and intersections using the joint *A*⋃*B, A*⋃*C, A*⋃*D, A*⋃*E* models. Multiple comparison correction was done using FDR with a threshold of *p* < 0.0*5*. Cerebellar data was resliced after the calculation of the unique partitions and the significance testing.

### Analysis of model weights

To assess similarity of semantic categories between cortex and cerebellum, the semantic space had to be broken into discrete categories instead of a smoothly continuous space. To do this the encoding model weights for the top 25% of voxels predicted by the word-level semantic model in each subject were concatenated together across subjects.

This was done separately in cortex and cerebellum, and then those were also concatenated together. Then the model weights were normalized across voxels and PCA was used to drop the number of dimensions from 985 to 86, which we chose because it explained 80% of the variance. These data were then clustered using spherical k-means into 5 clusters.

To choose the number of clusters we calculated inertia, which is the within-cluster sum of squares criterion, of the clustering algorithm for a range of clusters between 1 and 20 clusters. From this we calculated the point where the inertia changes from an exponential drop to a linear drop in inertia. This can also be defined as the point where the inertia is farthest from a linear line connecting the inertia at cluster 1 to the inertia at cluster 20. This point occurred at 5 clusters. (Supplemental Figure 11 shows the inertia across all clusters tested.)

To test for significance in category differences between cerebellum and cortex, a permutation test was done by shuffling voxels between the cortex and the cerebellum for each subject. The difference in the ratio of each category in the cerebellum as compared to the ratio of that category in the cortex was calculated for both the permutation set and the original data. The two-tailed p-value was calculated for each category as the ratio of the permutation difference greater than the absolute value of the original data difference plus the ratio of the permutation difference less than the negative absolute value of the original data. This was multiple comparison corrected using FDR with a threshold of *p* < 0.0*5*.

### Data Availability

A dataset including the data used in this study is being prepared for public release. Before the data are publicly available, they are also available upon request.

## Results

### Encoding model performance

To determine which aspects of language might be processed in the cerebellum, we created five feature spaces that span the hierarchy of language processing from sound to context-level meaning, including a spectral feature space, an articulatory space, a part-of-speech space, a word-level semantic space, and a context-level semantic space that combines information across words. Previous work has demonstrated that these feature spaces can capture these different components of language and predict BOLD responses in cortex(de Heer et al., 2017; Huth et al., 2016; Jain & Huth, 2018). We fit separate encoding models with each feature space using 5.4 hours of BOLD responses recorded while subjects listened to 26 different natural narrative stories taken from *The Moth Radio Hour*. Then, each model was used to predict responses to a *different* 10-minute story, and model performance was quantified as the correlation between the predicted and actual BOLD responses (R^2^). **Figure 2A** shows the prediction performance values for each feature space in one subject projected onto the SUIT cerebellar surface as well as prediction performance of each model in the cortex (similar maps for other subjects are in **Supplemental Figure 1**).

**Figure 2.**
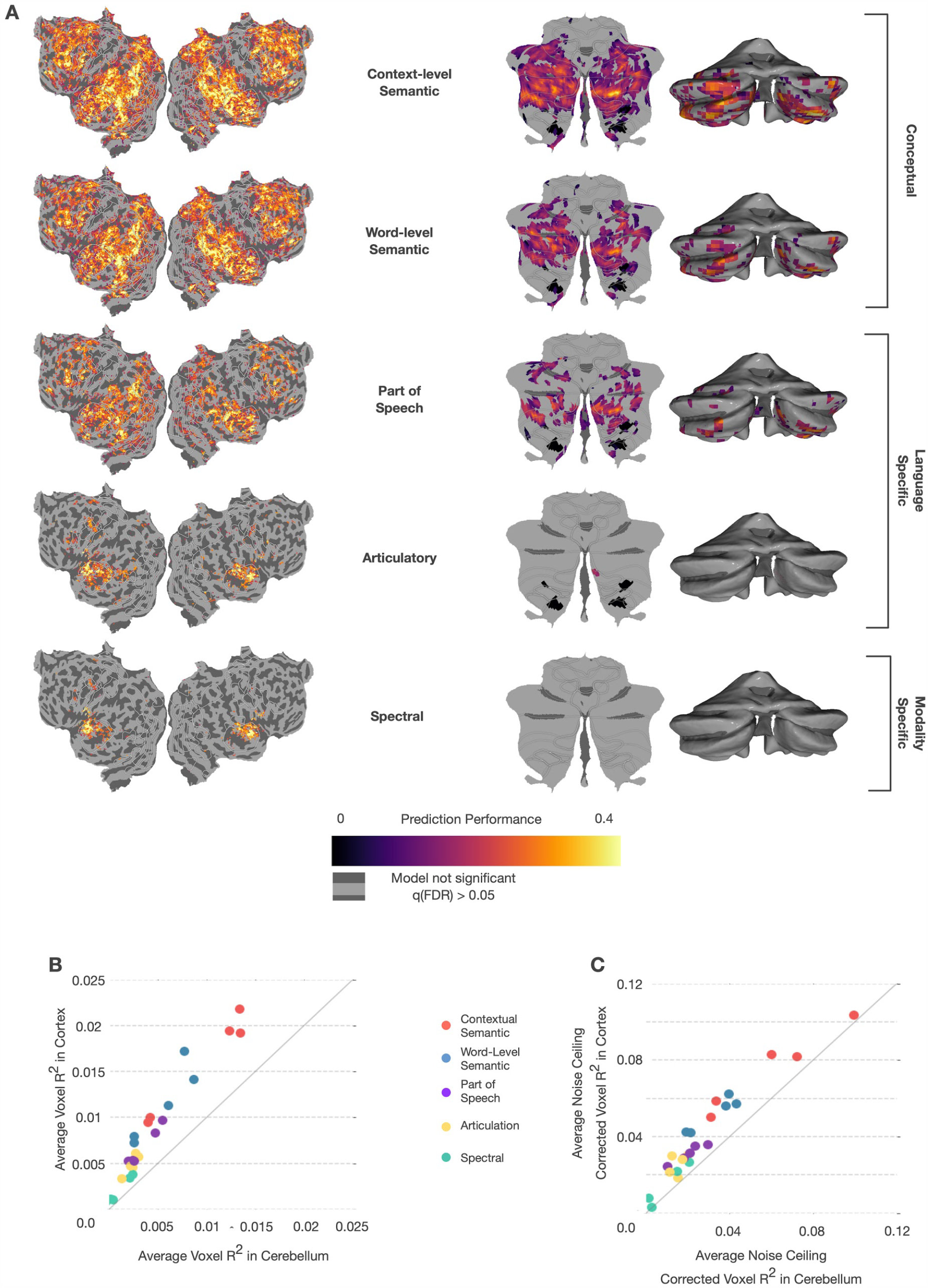
Prediction performance of encoding models based on five language feature spaces in cortex and cerebellum. Encoding models fit with 5.4 hours of BOLD data were tested against a held out story (10 minutes). (**A**) Correlation between predicted and actual BOLD response is plotted on flattened cortical and cerebellar surfaces for one subject (UT-S-02; other subjects are shown in **Supplementary Figure 1**). Significance testing for each model in each voxel was done using a one-sided FDR-corrected permutation test with a threshold of *p* < 0.0*5*. The higher-level models have better prediction performance in both cerebellum and cortex. In cortex, the areas best predicted by each of the three feature categories are spatially distinct. However, in the cerebellum, the areas best predicted by each feature space are highly overlapping. (**B**) To compare across subjects, we plotted average *R*^*2*^ across all voxels in the cerebellum and cortex for each subject and each feature space. The context-level semantic feature space has the highest predictive performance in both the cerebellum and cortex for all subjects. Performance scales roughly linearly in both cerebellum and cortex across the hierarchy of language representations, albeit with higher *R*^*!*^in cortex than cerebellum. (**C**) Because cortical and cerebellar BOLD responses might have different levels of noise, which could obscure differences in representation, we also computed noise ceiling-corrected correlations(Schoppe et al., 2016). This correction caused the average *R*^*2*^ to be less biased in favor of cortex (for corrected correlation flatmaps, see **Supplemental Figure 2)** and suggests that each feature space might be represented to a similar extent in cerebellum and cortex. However, overlapping prediction performance between different feature spaces in the cerebellum suggests that the cerebellum may not be separately representing each stage of language processing.

The spectral model uses a 256-dimensional, modality-specific feature space representing a mel-frequency spectrogram. This feature space is highly predictive of the primary auditory cortex along the transverse temporal gyrus. It does not significantly predict any voxel in the cerebellum (one-sided permutation test, q(FDR)<0.05), but it does appear to have diffuse low prediction performance across lobules VIIA, VIIB, and VIIIA. Of note, this is similar to previous results that showed cerebellar response to auditory stimulus along the medial portion of these lobules(Snider & Stowell, 1944). However, there appears to be no clustering of spectrally-selective voxels in the cerebellum, as is seen in the auditory cortex. This suggests that the cerebellum has no homologous area to the primary auditory cortex.

The articulatory model uses a 22-dimensional binary, language-specific feature space with each dimension representing one of the 22 articulations used in English (e.g. bilabial, back)(Levelt, 1993). In cortex, the articulatory space best predicts lateral, posterior temporal cortex along superior temporal gyrus. In the cerebellum, this feature space has diffuse prediction performance across lobules VIIIA and VIIIB, and significantly predicts a limited number of voxels in the medial posterior cerebellum (one-sided permutation test, q(FDR)<0.05). These areas are not traditionally considered motor speech areas(Callan et al., 2006; Manto et al., 2012) and thus this is unlikely to be due to covert rehearsal. This suggests that the cerebellum is not merely representing the articulations required to produce speech, and the lack of spatial clustering of well-predicted voxels further supports a lack of a homologous area to the auditory cortex.

The part-of-speech model uses a 17-dimensional binary, language-specific feature space, where each dimension represents one of 17 lexical classes (e.g. noun, verb, adjective). This feature space weakly but significantly predicts voxels covering a wide area of the cortex (one-sided permutation test, q(FDR)<0.05), including much of the frontal, temporal, and parietal lobes, with peak performance along the superior temporal lobe and near the intraparietal sulcus (IPS). In the cerebellum, this model significantly predicts voxels in many areas of the posterior lobe, with the highest model prediction performance in Crus I and II. This is a mid-level, language-specific feature space and its performance suggests that the cerebellum is largely representing information at a higher level than sound or articulations.

The word-level semantic model uses a 985-dimensional conceptual feature space that is based on word co-occurrence statistics across a large corpus of written English(de Heer et al., 2017; Deniz et al., 2019; Huth et al., 2016). This feature space captures semantic information under the assumption that *words that frequently occur in similar contexts carry similar meaning(FIRTH & R, 1957)*. The word-level semantic model predicts cortical voxels across regions in the frontal, parietal, and temporal lobes beyond core language-specific regions(Fedorenko et al., 2011, 2013). In the cerebellum, this model significantly predicts voxels in Crus I and II and lobules VIIIA and VIIIB. This conceptual model predicts much more response variance in the cerebellum and cortex than do lower-level models.

The best model in both the cerebellum and cortex is the context-level semantic model. This model builds on the word-level conceptual model by combining information across words. It uses the hidden state of a neural language model (LM) as a feature space. Neural LMs are artificial neural networks that learn to predict the next word in a sequence from past words. As a consequence, they learn a word’s meaning in context, improving upon the word-level model which is context-invariant(Lin et al., 2019; Radford et al., 2019; Tenney et al., 2019). Here, we used GPT(Jain & Huth, 2018; Radford et al., 2018), which is a popular neural LM. The feature space is 768-dimensional and the features are extracted from a middle layer of the LM that has previously been shown to be highly effective at predicting brain responses(Toneva & Wehbe, 2019). For each word, the past 25 words are used as context in the model. The context-level semantic model significantly predicts the largest number of voxels and most total variance across cortex, with peak prediction performance in frontal, parietal and temporal cortex. In the cerebellum, this model yields very high prediction performance across most of the posterior cerebellum, including Crus I and II and lobules VIIIA and VIIIB.

To compare model performance between the cerebellum and cortex directly, we computed the average performance of each model in the cerebellum and cortex for each subject. **Figure 2B** shows that there is a linear relationship between model performance in the cerebellum and cortex, suggesting that language might be represented similarly in these two structures. To account for the possibility that BOLD signal-to-noise varies systematically between cortex and cerebellum, we also adjusted the estimated correlation for each voxel using a standard technique(Schoppe et al., 2016). **Figure 2C** shows these results when accounting for the difference in signal-to-noise variance between cortex and cerebellum. Here, the pattern of results is largely the same, but prediction performance in the cerebellum is more similar to that of cortex. In both cases however, cerebellar voxels that are well predicted by each feature space are highly overlapping. This could be caused by the feature spaces carrying overlapping information with each other, making it difficult to interpret the results from each feature space independently. To disentangle these representations and explore the differences between cortex and cerebellum in more detail, we next performed a variance partitioning analysis.

### Variance Partitioning

The previous model comparison found that many voxels in the cerebellum can be significantly predicted by multiple feature spaces. These voxels might genuinely represent information from multiple feature spaces. Indeed, the increased neuronal density of cerebellum compared to cortex(Herculano-Houzel, 2010) raises the chance that individual cerebellar voxels contain information from multiple feature spaces. However, this effect could also be a consequence of correlations, or shared information, between the feature spaces. To disentangle possible overlaps in information across the five feature spaces within each voxel we used variance partitioning, a statistical technique for determining how much variance can be uniquely explained by each set of features(de Heer et al., 2017; Lescroart et al., 2015). This enables us to distinguish between overlapping but distinct representations and seemingly overlapping representations that actually reflect correlations between features. For example, variance partitioning would allow us to disentangle if say 50% of the voxel responds to conceptual information and another 50% to auditory information, or if 100% of the voxel response is to some feature that is correlated with both auditory and conceptual information.

Our first variance partitioning analysis shows how much variance each feature space uniquely explains above all other feature spaces for each voxel, and the second shows how much overlap there is between each feature space and the context-level semantic feature space.

### Unique Variance Explained

The results in **Figure 2a** showed negligible, localized prediction performance of low-level models in the cerebellum, suggesting that little low-level language processing was occurring there. However, that result did not account for the possibility that higher-level feature spaces could also capture some low-level information. To test for this, we used variance partitioning to find the unique variance explained of each feature space in order to test whether the lower level models have any unique contribution to representations in the cerebellum. This was done by first fitting a five-way union encoding model with a concatenation of all the feature spaces. Variance explained by any of the five feature spaces should be explained by this five-way union model. Then we fit five additional encoding models, each combining four of the five feature spaces. Each of these models should explain all the variance captured by the five-way union model except for variance that is uniquely explained by the feature space that was left out. To estimate the unique variance explained by each feature space, we then subtracted the variance explained in the four-way model excluding that feature space from the five-way union model (for additional details, see **Methods**). We also used these models to estimate the size of the non-unique partition, which contains any variance that can be explained by more than one of the five feature spaces. The size of this partition was calculated by subtracting each of the unique variance partitions from the five-way union model. For this analysis, we only considered voxels that were significantly predicted by the five-way union model (one-sided permutation test, q(FDR)<0.05).

**Figure 3A** shows the unique variance explained by each feature space as well the non-unique partition for each voxel in the cerebellum and cortex projected onto the flattened surface for one subject (other subjects can be seen in **Supplemental Figure 3**). The non-unique partition is the largest partition overall, suggesting that much of the variance explained by these feature spaces cannot be specifically allocated to one feature space. It is important to note that this category includes all possible combinations of the feature spaces and does not mean that the variance is explained equally well by each of the five feature spaces. Among the unique partitions, both the context-level semantic model and spectral feature space explain variance significantly greater than zero (one-sided permutation test, q(FDR)<0.05). However, the spectral model explains significantly (two-sided permutation test, q(FDR)<0.05) more variance in cortex than in the cerebellum. **Figure 3B** shows the unique variance explained for each feature space averaged across voxels for all subjects (only including voxels that were significantly predicted by the union model). We compared mean partial correlations 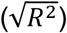 between cerebellum and cortex for each partition using a permutation test. The result shows that the spectral feature space and non-unique partitions explain significantly less variance in the cerebellum than in cortex. When correcting for differences in signal-to-noise (**Supplemental Figures 4 & 5**), the context-level semantic and word-level semantic feature spaces uniquely explain significantly more variance in the cerebellum than cortex, and the spectral feature space uniquely explains significantly less.

**Figure 3.**
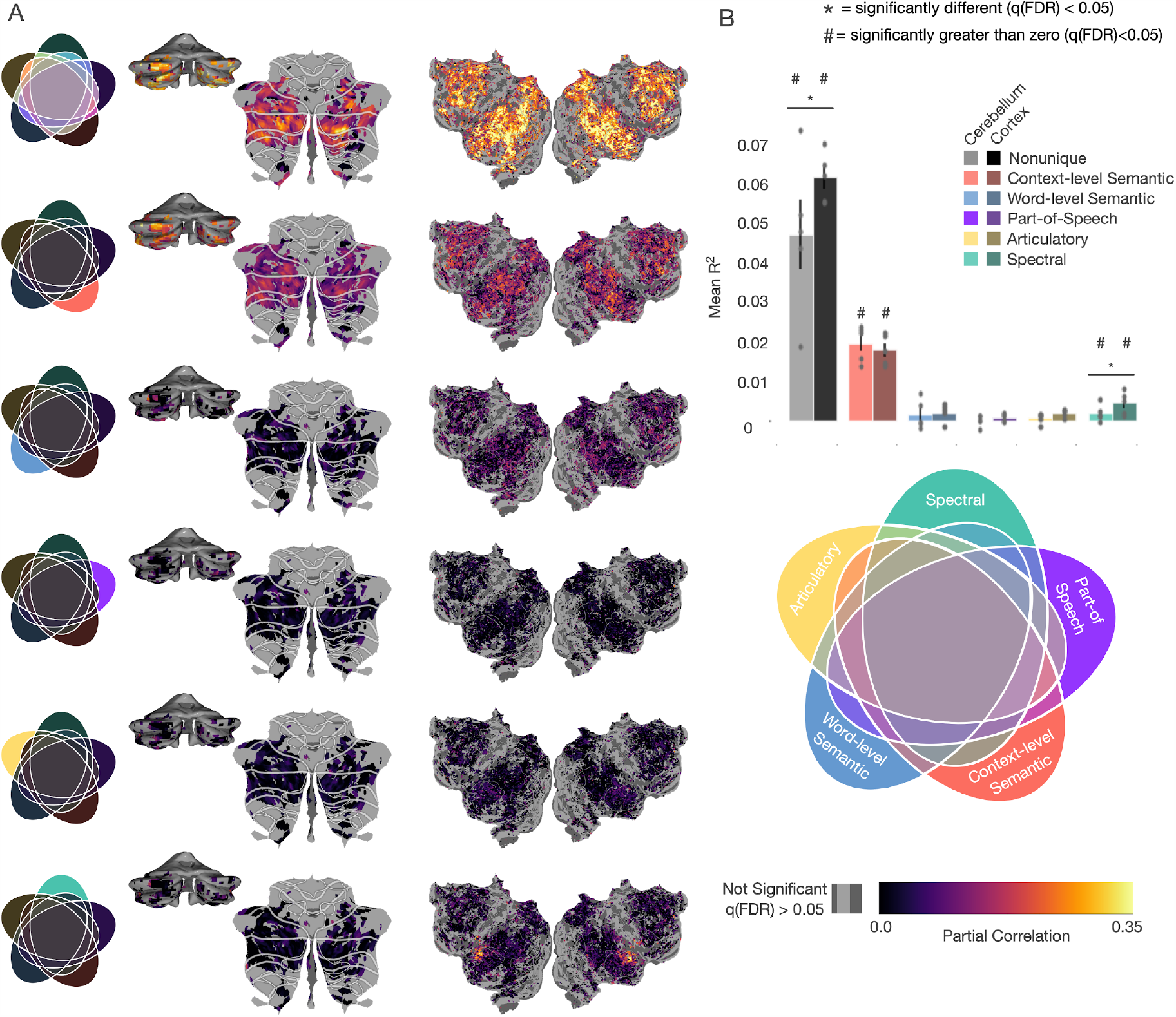
Unique variance explained by each feature space. To determine how much variance is uniquely explained by each feature space, six new encoding models were fit: a union model containing a concatenation of all feature spaces, and five encoding models each containing a concatenation of four of the five feature spaces. The unique contribution of each feature space was then determined by subtracting the variance explained by the four-way concatenation model without that feature space from the union model. This shows how much variance can be explained by each feature space above and beyond the other four. Additionally, the amount of non-unique variance—i.e., any that can be explained by more than one feature space—was determined by subtracting the 5 unique variances from the union. (**A**) The voxelwise partial correlation 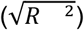 for each feature space for subject UT-S-02, projected onto the cortical and cerebellar surfaces (similar maps for other subjects are in **Supplemental Figure 3)**. Only voxels that were significantly predicted (one-sided permutation test, q(FDR)<0.05) by the 5-way union model are displayed. (**B**) Mean correlations for significant voxels in the cerebellum and cortex across all subjects. The non-unique partition contains the most variance in both cortex (darker) and cerebellum (lighter), but is significantly smaller in cerebellum than cortex (two-sided permutation test, q(FDR)<0.05). The modality-specific spectral feature space explains significantly less variance in the cerebellum as compared to the cortex. Additionally, the modality-specific feature spaces do not uniquely explain any significant variance (one-sided permutation test, q(FDR)<0.05), while the context-level semantic space uniquely explains the most variance. This further supports the hypothesis that the cerebellum is largely representing language at a high, conceptual level.

**Figure 4.**
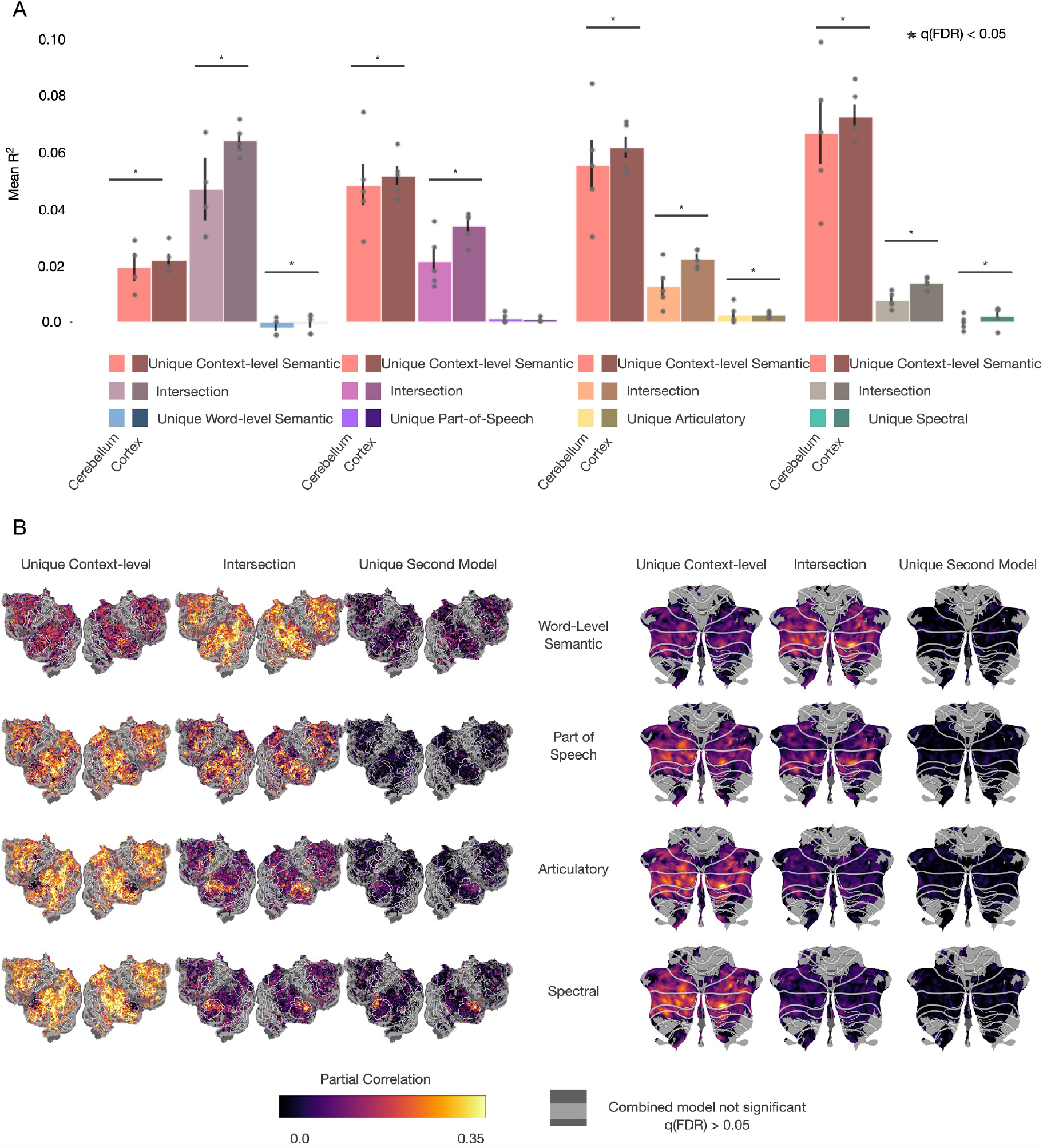
Variance partitioning between the context-level semantic feature space and each of the other feature spaces. To quantify the amount of overlap between the context-level semantic feature space and each of the four other feature spaces, three models were fit for each pair of feature spaces, including the concatenation of both feature spaces and each feature space individually. (**A**) For each pair feature spaces, the variance uniquely explained by the context-level feature space, that uniquely explained by the second feature space, and the intersection between the two is compared between the cerebellum and cortex, averaged over all subjects. The intersection—variance that could be explained by either feature space—for every pair is smaller in the cerebellum than in the cortex (two-sided permutation test, q(FDR)<0.05). Additionally, the unique partition for the spectral feature space is significantly smaller in the cerebellum than in cortex. This shows that the high prediction performance of the context-level semantic feature space in cerebellum is not merely due to correlations with modality- and language-specific information. Instead, the context-level features uniquely explain a large amount of variance that the other features cannot. (**B**) For each pair of models, the variance in each partition in each voxel was projected onto cortical and cerebellar flatmaps (**Supplemental Figures 8, 9, & 10**, noise-ceiling corrected version to account for differences in signal-to-noise across the brain). Only voxels that were significantly predicted by each union model (one-sided permutation test, q(FDR)<0.05) are shown. There is substantially lower variance explained by the intersection between the context-level semantic feature space and the language and modality-specific feature spaces in the cerebellum than in cortex. Additionally, the unique contributions for these feature spaces in the cerebellum is near zero and is not spatially localized. This lack of spatial localization further suggests that there is no hierarchy of language processing in the cerebellum, and these results provide strong support for the hypothesis that the cerebellum only represents high level, conceptual features of language, rather than low-level features.

Both of these results suggest that the cerebellum is primarily representing language at a conceptual level and that these results are not simply due to neuronal pooling within voxels or shared representations. However, the fact that the largest proportion of variance is in the non-unique partition means that this analysis alone can not rule out the possibility for low-level language representations in the cerebellum.

### Pairwise partitioning

In the first variance partitioning analysis, we found that the context-level semantic feature space explains the most unique variance explained and that the spectral model explains significantly less variance in the cerebellum than in cortex. This suggests that the cerebellum may not be representing information at modality and language-specific levels. However, the largest partition in both the cerebellum and cortex was the non-unique partition, which contains variance that could be explained by more than one feature space. Thus, that analysis alone cannot rule out the possibility that low-level features are represented in cerebellum. To test the hypothesis that the cerebellum is exclusively representing language at a conceptual level, we performed a second variance partitioning analysis where each feature space was separately compared to the context-level semantic feature space. We fit four union models by concatenating the context-level semantic features with each one of the four other feature spaces. The variance explained by each union model was then compared to models fit with each feature space individually in order to determine both the unique contribution of each feature space and the size of their intersection. For each pair of feature spaces, analyses were restricted to voxels that were significantly predicted by the union model. If the cerebellum was only representing information at the conceptual level, we would expect to find low unique variance explained by the modality- and language-specific feature spaces and a high shared intersection with the word-level semantic feature space.

The results of this pairwise variance partitioning analysis replicate previous results(de Heer et al., 2017) showing that in cortex there is a unique contribution of both the spectral and articulatory feature spaces in different cortical areas. However, this does not appear to be true in the cerebellum. **Figure 4** shows the results of pairwise variance partitioning between the context-level semantic feature space and each of the other four feature spaces. **Figure 4A** shows the mean partial correlation for each pair of feature spaces in both the cerebellum and cortex across voxels and subjects. The variance explained by the intersection of each pair of models is significantly less in the cerebellum than in cortex (two-sided permutation test, q(FDR)<0.05). This shows that the information present in the lower level feature spaces contributes less to the explainable variance in the cerebellum and supports the hypothesis that the cerebellum is primarily representing high level, conceptual information. Additionally, the unique contribution from the modality- and language-specific feature spaces are negligible; the spectral feature space explains significantly less variance in the cerebellum, while the articulatory feature spaces explain significantly more variance in the cerebellum, although this partition is small in both the cerebellum and cortex. When accounting for differences in signal-to-noise (**Supplemental Figures 8, 9, & 10)**, all of the unique contributions from the secondary models become significantly less in the cerebellum than in cortex. Additionally, the differences in the intersections between the cerebellum and cortex are no longer significant. However, the unique contribution from the context-level semantic feature space is significantly larger in all cases in the cerebellum and cortex. While the noise-ceiling corrected results are different due to the differences in BOLD signal in the cerebellum as compared to cortex, the significantly larger variance explained by the context-level semantic feature space in the cerebellum still supports the hypothesis that the cerebellum is uniquely representing highly conceptual semantic information. **Figure 4B** shows these results for one subject projected onto the corresponding cortical and cerebellar surfaces (cortical maps for all other subjects can be found in **Supplemental Figures 6** and **7**). Only voxels that were significantly predicted by the union model (one-sided permutation test, q(FDR)<0.05) are displayed.

Very little variance in the cerebellum is explained uniquely by any features other than the context-level semantic space. In both the cerebellum and cortex, there is a high amount of variance explained by the context-level semantic feature space and in the intersection with the word-level semantic feature space. This is not surprising, given that the context-level semantic space has the highest predictive performance of any of the feature spaces and that the word-level and context-level semantic spaces contain related semantic information. However, there is very little overlap of variance explained between the context-level semantic feature space and the three modality- and language-specific feature spaces. This demonstrates that the high performance of the conceptual feature spaces is not merely due to this feature space being correlated with low-level information. The negligible unique contribution of the modality- and language-specific features in the cerebellum further supports the hypothesis that the cerebellum is primarily representing conceptual representations. Finally, any variance explained by the modality- and language-specific feature spaces is not anatomically localized within cerebellum, which suggests that the cerebellum does not contain localized low-level language processing areas. The reduced representation of language specific feature spaces in the cerebellum further suggests that the cerebellum does not participate in language processing per se, but supports cognition more generally.

### Semantic selectivity within the cerebellum

Our results thus far suggest that the cerebellum is not involved with language-specific processing, as there is little or no unique variance explained in cerebellum by the part-of-speech, articulatory, or spectral feature spaces. Instead, language representations in the cerebellum appear to be dominated by conceptual semantic features. Yet all semantic representations are not alike: in cortex, earlier work revealed a patchwork tiling of areas that represent different semantic categories across much of prefrontal, parietal, and temporal cortex(Jeffrey R. Binder et al., 2009; Huth et al., 2016; Noppeney & Price, 2004). It is possible that the cerebellum represents a different range of semantic categories than cortex, and it seems likely that different categories are represented in distinct areas within the cerebellum. Following the procedure detailed in Huth et al. (2016), we used the word-level semantic feature space to analyze and interpret the model weights and thus reveal the semantic selectivity of each voxel in the cerebellum. Due to the lack of tools currently available for interpreting context-level semantic models, we chose to use the word-level model, which also explains a large proportion of response variance in the cerebellum.

To demonstrate how encoding models can be analyzed per voxel, **Figure 5A** shows the word level semantic regression weights projected into a three-dimensional semantic space that was previously constructed from a group of subjects using principal components analysis (PCA)(Huth et al., 2012). This lower-dimensional space is purely used for visualization purposes. Here projections on the first, second, and third principal components (PCs) are mapped into the red, green, and blue color channels respectively, for each voxel and then projected onto the SUIT cerebellar surface. The color wheel shows approximately which semantic category each color on the maps represents. **Figure 5A** shows the posterior view of one subject’s (UT-S-02) cerebellum as well as the flattened cerebellar surface in SUIT space. Within the SUIT space, functional regions of interest are mapped out which include anterior foot (AF), hand(AH), and mouth (AM); posterior foot(PF), hand(PH) and mouth (PM); and anterior and posterior eye movement areas (AE and PE) that are active during saccades. A histogram of correlations for all voxels in subject UT-S-02 is shown in **Figure 5B**. This histogram shows a distribution with a long tail, with the example well-predicted voxel (voxel 1685) marked in blue.

**Figure 5.**
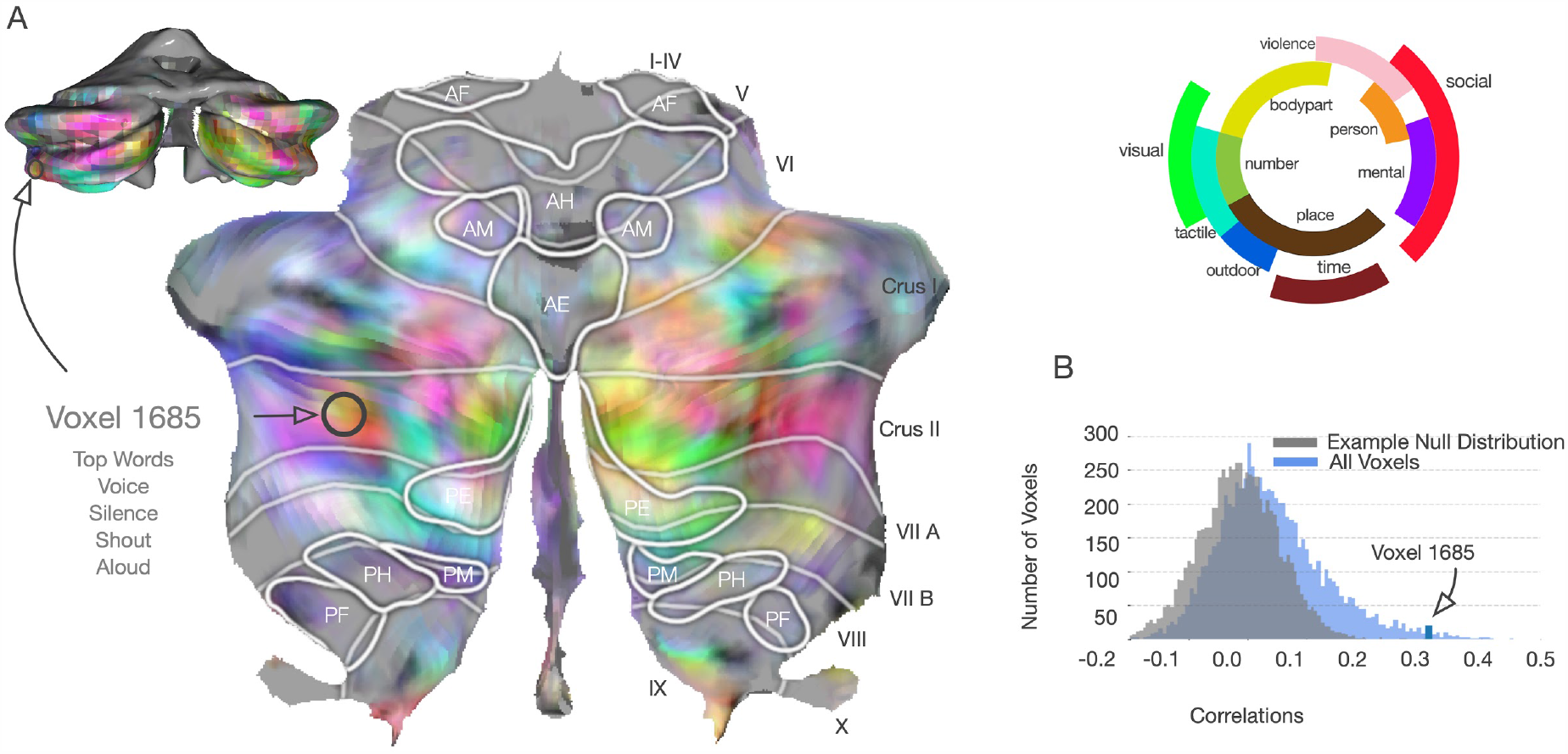
Word-level semantic model weight interpretation. Post hoc analysis of encoding models enables us to interpret what type of semantic information is represented in each voxel. Here we used the word-level semantic feature space to interpret one individual voxel and to broadly map semantic representations across the cerebellum. (While the context-level semantic space is more predictive, we lack tools for interpreting its representations.) In the word-level space, encoding models predict the response of each voxel to each word. We used the model to find words with the largest predicted response in one voxel (voxel 1685 in subject UT-S-02), which were “voice”, “silence”, and “shout”, suggesting that this voxel represents concepts related to social communication. To visualize representations across many voxels, we reduced the encoding model weights to three dimensions by projecting them onto a low-dimensional semantic space identified in a previous experiment(Huth et al., 2016), and then mapping these projections to RGB color channels. (**A**) The RGB values for each voxel are projected onto the SUIT cerebellar surface for subject UT-S-02. Different colors correspond to selectivity for different concepts in the semantic space (illustrated by the legend, right). This map suggests that the cerebellum contains representations of many different concepts. This histogram (**B**) shows the range of correlations for each voxel in this subject, with the example voxel marked in blue, and null distribution in gray.

**Figure 6.**
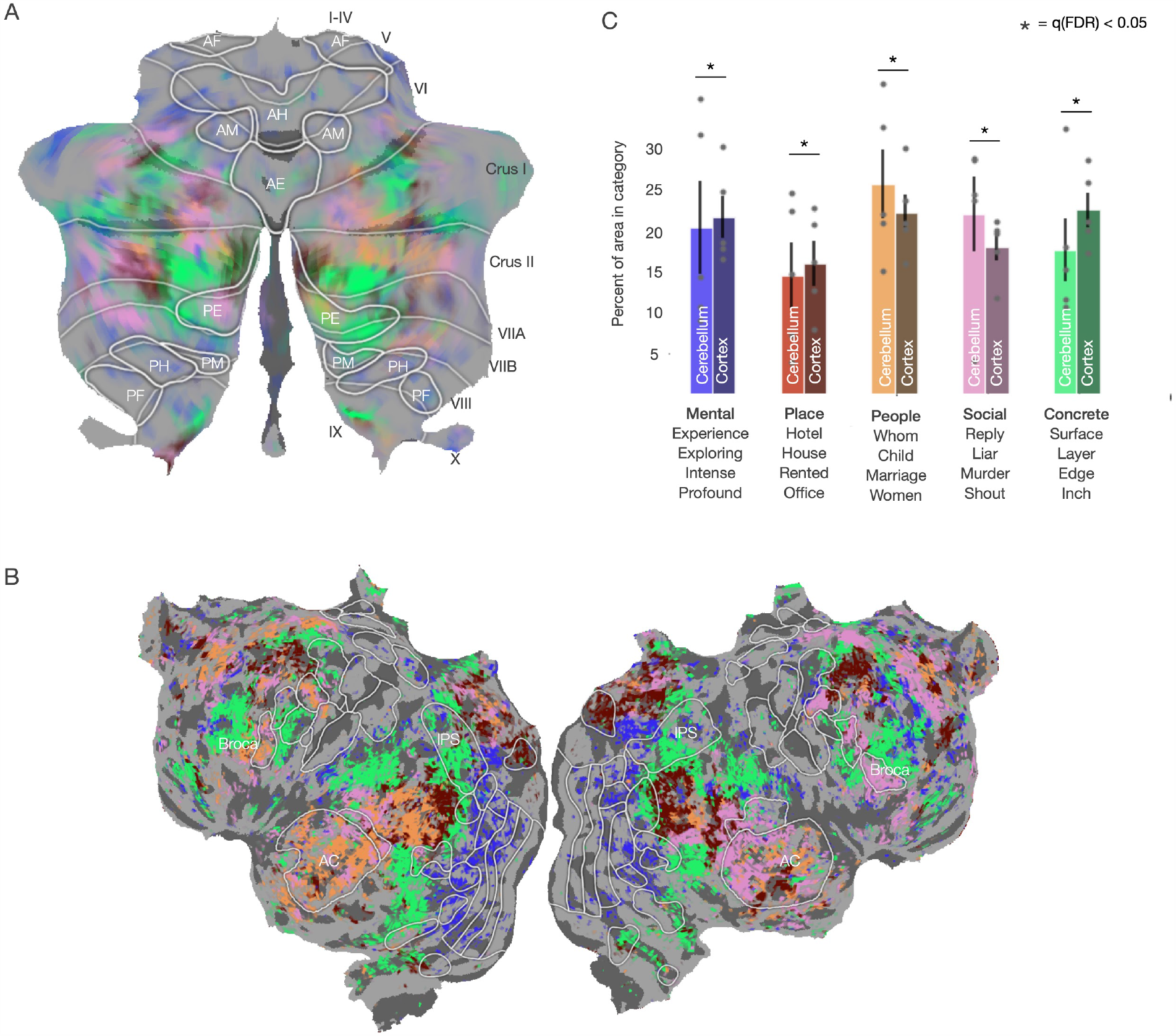
Differences in semantic representations between cerebellum and cortex. To check for differences in semantic representations between the cerebellum and cortex, word-level encoding model weights from both cerebellum and cortex in all subjects were concatenated, including only the top 20% best-predicted voxels. This matrix was then clustered using spherical k-means into 5 clusters, which fell at the inflection point in the inertia graph (**Supplemental Figure 11**). For visualization, the centroid for each cluster was transformed into the same RGB space used in **Figure 5**, and each voxel in that cluster was assigned that color. (**A**) shows the cluster distribution for one subject, UT-S-02, across the cerebellum and (**B**) cortex. Voxels falling into each cluster are found in both the cerebellum and cortex in every subject. (**C**) To test for differences in representation between cortex and cerebellum, the percentage of cortical and cerebellar voxels in each cluster were compared across all subjects. Each cluster was named qualitatively according to the most similar words to the cluster centroid. The four most similar words to each cluster centroid are listed below the label name. Significantly more voxels in the cerebellum were highly responsive to social categories (two-sided permutation test, q(FDR)<0.05), i.e. the “social” and “people” clusters, than in cortex. Conversely, significantly fewer voxels in the cerebellum were responsive to the “mental”, “concrete”, or “place” clusters than in the cortex. This shows that the cerebellum is largely representing the same semantic categories as cortex, but that there is a slight bias towards social categories.

Additionally, **Figure 5A** lists the five words that the word-level semantic encoding model predicts will elicit the largest response in this example voxel, which are “voice”, “silence”, “shout”, and “aloud”. These words were found by taking the dot product of the voxel weight vector with the word-level semantic feature matrix (see **Methods** for details). This voxel seems selective for concepts related to social communication and sound. Similar analysis could be performed for each voxel, but would be large and difficult to interpret. However, by representing semantic weights as a color we can better understand large-scale patterns of semantic information. For example, Crus I and Crus II seem to be selective for many different semantic categories, such as *social* and *violence* which can be found in medial Crus I.

### Comparing semantic representations between cerebellum and cortex

The semantic map in **Figure 5A** shows that different areas in the cerebellum represent different categories of words. Yet it is not clear from this map whether semantic representations in the cerebellum are similar to those found in cortex. To quantify the semantic categories represented in the cerebellum and cortex, we compared the fraction of voxels that represent different semantic categories using a cluster analysis. We concatenated model weights for the top 20% best predicted voxels in cerebellum and cortex from each subject, then clustered the voxels into 5 discrete categories using spherical k-means clustering (5 clusters was the elbow point of the inertia curve, see **Supplemental Figure 11**; similar results are also obtained with different numbers of clusters). **Figure 6A** and **B** show cerebellar and cortical flatmaps with the clustered voxels colored according to their assigned cluster in one subject (similar maps for other subjects can be found in **Supplemental Figure 12**). The label for each cluster was determined qualitatively from the most similar words to each cluster centroid (**Figure 6C** lists the clusters, their top words, and their assigned label).

Voxels belonging to every semantic cluster were found in both the cerebellum and cortex. **Figure 6C** shows the percentage of cerebellar voxels in each cluster as compared to the percentage of cortical voxels in each cluster, averaged across subjects. The category with the highest percentage of voxels in the cerebellum is the “people” category and the category with the lowest percentage is the “place” category.

Because voxels in all clusters are found in both the cerebellum and cortex, it is possible that the cerebellum is receiving input from all areas of cortex. If this was true, we would expect to find an equal percentage of well-predicted cerebellar voxels in each cluster as there are in cortex. However, all clusters had significantly different percentages of voxels in the cerebellum as compared to cortex (two-sided permutation test, q(FDR)<0.05). The “social” and “people” clusters have a higher percentage of voxels in the cerebellum than in cortex, and the “mental”, “concrete”, and “place” clusters have a lower percentage of voxels in cerebellum than in cortex. This suggests that there is not a one-to-one mapping from cortex to the cerebellum and that the cerebellum is more responsive to social semantic information.

## Discussion

This study examined how language is represented in the human cerebellum. Using voxelwise encoding models trained within each subject using large amounts of fMRI data, we found that high-level language feature spaces–context-level and word-level semantics–were better able to predict cerebellar BOLD responses than low-level language feature spaces, such as part-of-speech and articulations. Additionally, the low-level feature spaces do not uniquely predict any voxel in the cerebellum above the context-level semantic model, which is not true in the cortex. Lastly, using the model weights from the word-level semantic model, we found that there is an overrepresentation of social and people semantic categories in the cerebellum as compared to cortex. These results suggest that the cerebellum is (1) representing language at a conceptual level, and not at modality- or language-specific levels, (2) that there is not a homologous area to auditory cortex in the cerebellum, and (3) that the cerebellum is more responsive to social semantic components of language than cortex. As has been seen previously(King et al., 2019), there does not appear to be any functional relevance to lobule boundaries as we do not observe any pattern of language processing that corresponds to the lobule boundaries.

One complication in interpreting the results of this study is due to the use of BOLD fMRI, in particular in relation to the cerebellum. The cerebellum has a significantly different metabolic demand than cortex(Vaishnavi et al., 2010) due to its cellular architecture. This changes the demand for oxygenated blood and thus the BOLD signal. It has previously been demonstrated that only activity in granule cells and mossy fibers affects the BOLD signal(Caesar et al., 2003; Mathiesen et al., 2000) in the cerebellum, but not activity in the Purkinje cells which are the sole output from the cerebellum to cortex. This implies that our models do not include representations of what the cerebellum is outputting back to the cortex and thus may not directly address the computation the cerebellum is performing. However, the input to the cerebellum– granule cells and mossy fibers– is still an important half of the equation and this work furthers our understanding of what kinds of representations are being sent to the cerebellum.

One point of contention with our methodology is using a natural stimulus. While natural stimuli can make interpretation of the results more difficult, it is a much richer stimulus set for analysis. Additionally, it is less biased than other experimental methods that preselect a small number of categories or stimuli. And while we have few subjects, we have collected a large amount of data per subject. This amount of data allows for the regression models to theoretically account for most stimulus correlations. Additionally, by using a prediction methodology we are able to compute the variance explained by each feature space which allows us to quantify how well each model does at prediction which few other methods allow for.

Many theories exist for how the cerebellum represents cognitive information based on the uniformity of its cellular architecture. This architecture is believed to suggest that the cerebellum is performing a similar function throughout the structure. Additionally, the cerebellum has long been considered a major region in motor response and motor learning(Manto et al., 2012). Yet since the 1980s, the cerebellum has been known to reliably respond during cognitive tasks(Leiner et al., 1986) such as language processing(Petersen et al., 1988) and that lesions to the posterior lobe of the cerebellum results in language deficits(Schmahmann & Sherman, 1998). The fact that both fine motor control processing and cognitive processing elicit strong responses from the same architecture has long been considered a contradiction. In an effort to reconcile the cerebellum as both a cognitive area and a motor area, previous reports have speculated that the cerebellum is involved in some low-level component of cognitive tasks such as low-level auditory processing(Petacchi et al., 2005) or motor planning in speech(Jürgens, 2002).

Surprisingly, our results show that low level spectral and articulatory feature spaces do not uniquely predict any area of the cerebellum better than high-level feature spaces. However, the spectral and articulatory models best predict areas around auditory cortex and the STG. This shows that these feature spaces successfully capture auditory information and that this information does not appear to be present in the cerebellum. The cerebellum is very likely involved with motor components of speech production. However, we found no evidence of receptive articulatory representations in the cerebellum, suggesting it must be involved in more than just motor components of speech perception. Our results imply that the cerebellum lacks any form of localized low-level auditory processing area and that the role of the cerebellum in language processing is at a higher level than previously thought.

An important future step is to clarify the relationship between language representations in the cerebellum and existing theories of cerebellar function. The universal cerebellar transform (UCT) theory is the predominant theory of cerebellar computation(Diedrichsen et al., 2019), positing that the cerebellum performs a single computation across all tasks, both cognitive and motor. A commonly proposed computation is prediction error(Kawato & Gomi, 1992; Mariën & Manto, 2018). One way to look at whether the cerebellum is involved in prediction is through surprisal which is a measure of the probability of a word occurring in a sentence given the previous word. Thus a word with a high surprisal is likely to have a high prediction error. Since the context-level semantic model is using a neural network based language model, it inherently captures some elements of surprisal(Berger et al., 1996). However, the context-level semantic model best predicts both the cerebellum and cortex which suggests that the cerebellum is not uniquely computing surprisal.

In language processing, many processes are specific to auditory communication such as the spectral and articulatory features spaces. However, the higher order semantic features seem to be more broadly used by the default mode network. Given that the cerebellum does not appear to be involved in the lower level language processing, this implies that the cerebellum is not participating in language processing per se, and is likely only involved in cognitive processing. This theory could explain many of the language deficits seen in patients with CCAS and autism. Both of these disorders are associated with cerebellar damage or morphological changes and both often see deficits in language processing(Stoodley & Schmahmann, 2009). However, the deficits are not specifically related to speech production or the ability to interpret sound into phonemes and words, which are low-level language-specific processes. Rather, the language deficits in CCAS and autism more often present as conceptual deficits, with a loss of understanding of fine-tuned semantic specificity and social dynamics(Kelley et al., 2006), such as understanding sarcasm and non-explicit language. Much like the cerebellum being involved in the fine-tuning of motor commands over a continuous three-dimensional space, it is possible that the cerebellum is similarly involved in the fine-tuning of a conceptual cognitive space.

## Supplemental Figures

**Supplemental Figure 1.**
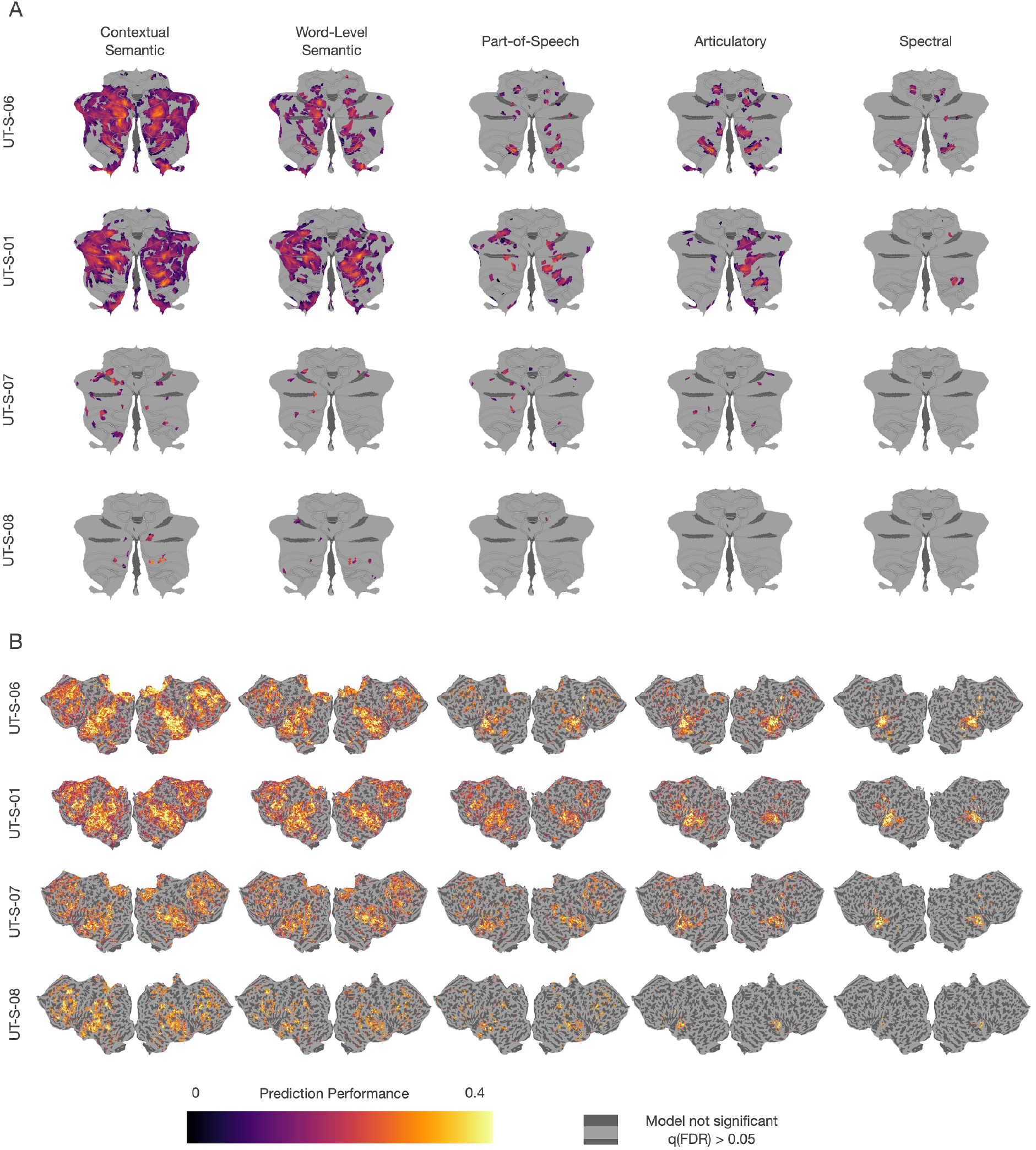
Prediction performance of encoding models based on five language feature spaces in cortex and cerebellum. Encoding models fit with 5.4 hours of BOLD data were tested against a held out story (10 minutes). The correlation between predicted and actual BOLD response is plotted on flattened cerebellar (**A**) and cortical (**B**) surfaces for each subject. Significance testing for each model was done using a one-sided FDR-corrected permutation test with a threshold of *p* < 0.0*5*. The higher level models have better prediction performance in both cerebellum and cortex. In cortex, the areas best predicted by each of the three categories of feature spaces are spatially distinct. However, in the cerebellum the areas best predicted by each of the feature spaces is highly overlapping. This suggests that there is a hierarchy of language processing in cortex but not in the cerebellum.

**Supplemental Figure 2.**
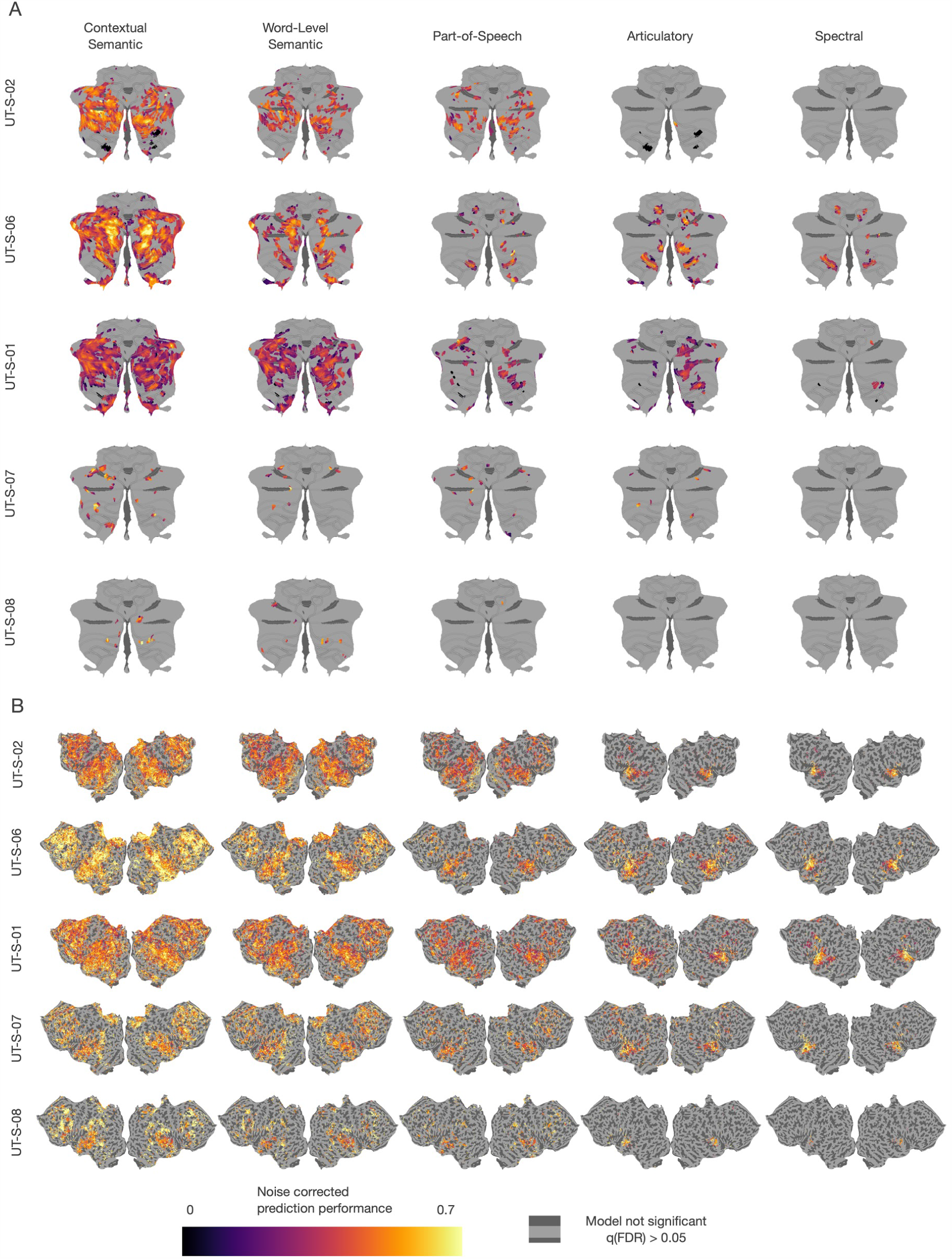
Prediction performance of encoding models based on five language feature spaces correcting for difference in signal-to-noise in cerebellum and cortex. Encoding models fit with 5.4 hours of BOLD data were tested against a held out story (10 minutes). The correlation between predicted and actual BOLD response is plotted on flattened cerebellar (**A**) and cortical (**B**) surfaces for one subject (UT-S-02).The correlations were noise-ceiling corrected using standard techniques to account for differences in BOLD signal-to-noise ratio in the cerebellum and cortex. Significance testing for each model was done using a one-sided FDR-corrected permutation test with a threshold of *p* < 0.0*5*. The higher level models have better prediction performance in both cerebellum and cortex. In cortex, the areas best predicted by each of the three categories of feature spaces are spatially distinct. However, in the cerebellum the areas best predicted by each of the feature spaces is highly overlapping. This suggests that there is a hierarchy of language processing in the cortex and not in the cerebellum

**Supplemental Figure 3.**
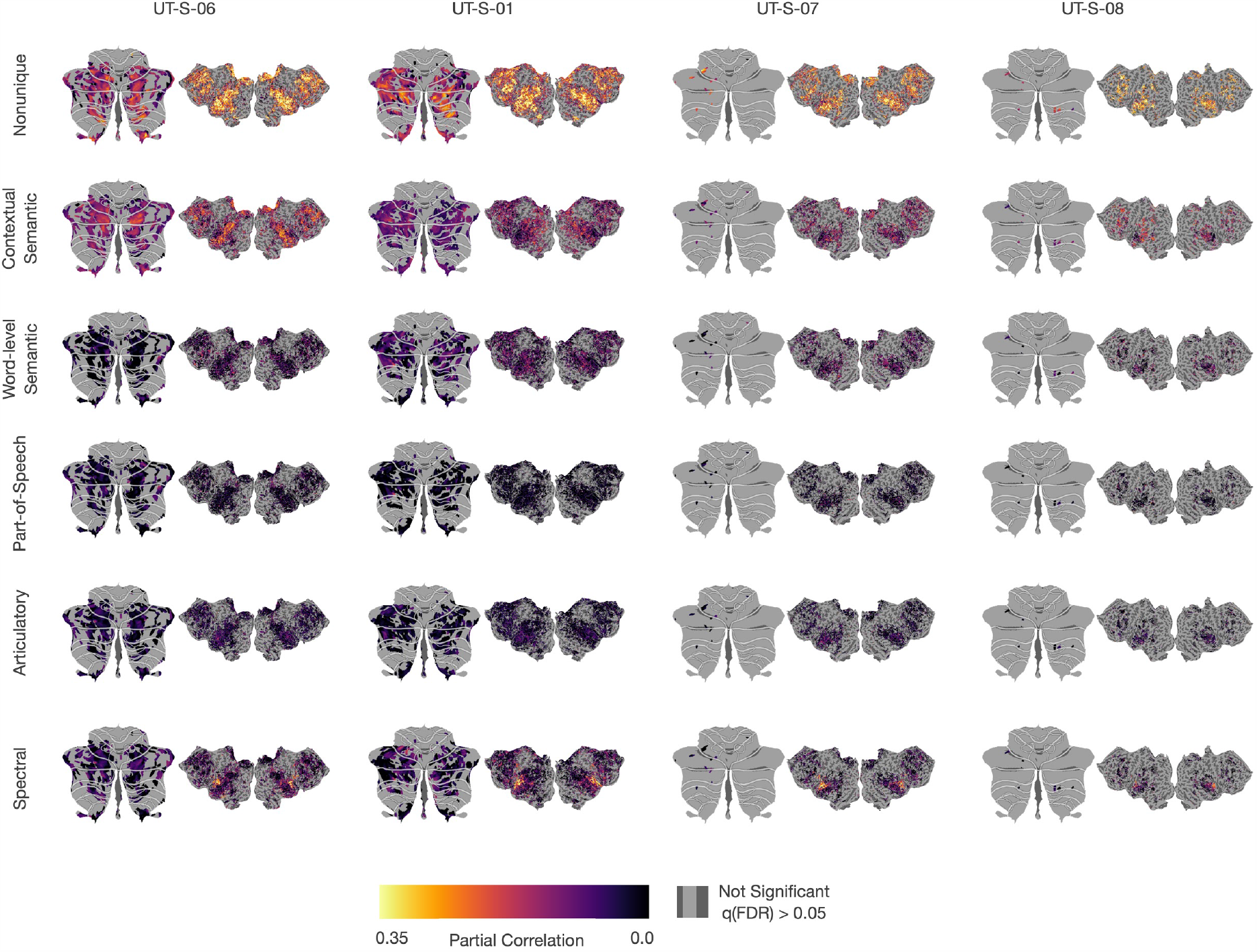
Unique variance explained for each feature space. To determine the unique variance explained by each feature space, a union encoding model was fit with a concatenation of all feature spaces in addition to five other encoding models - each a concatenation of four of the feature spaces. The unique contribution of a model can be determined by the subtraction of the four way concatenation model without that feature space from the five way union model. Additionally, the amount of overlap between the feature spaces can be characterized by the non unique partition. (**A**) The unique variance explained by each feature space in each additional subject were projected onto the cortical and cerebellar surfaces. Only significant voxels (one-sided permutation test, q(FDR)<0.05) from the 5-way joint model are displayed. The partition that explains the most variance in both cortex and cerebellum is the nonunique partition, which shows that there is overlap in the information in the feature spaces. The largest unique partition in the cerebellum and cortex is the context-level semantic feature spaces which shows that both areas likely represent conceptual information. The largest difference in unique partitions between the cerebellum and cortex is the spectral feature space, which does not uniquely explain any region of the cerebellum.

**Supplemental Figure 4.**
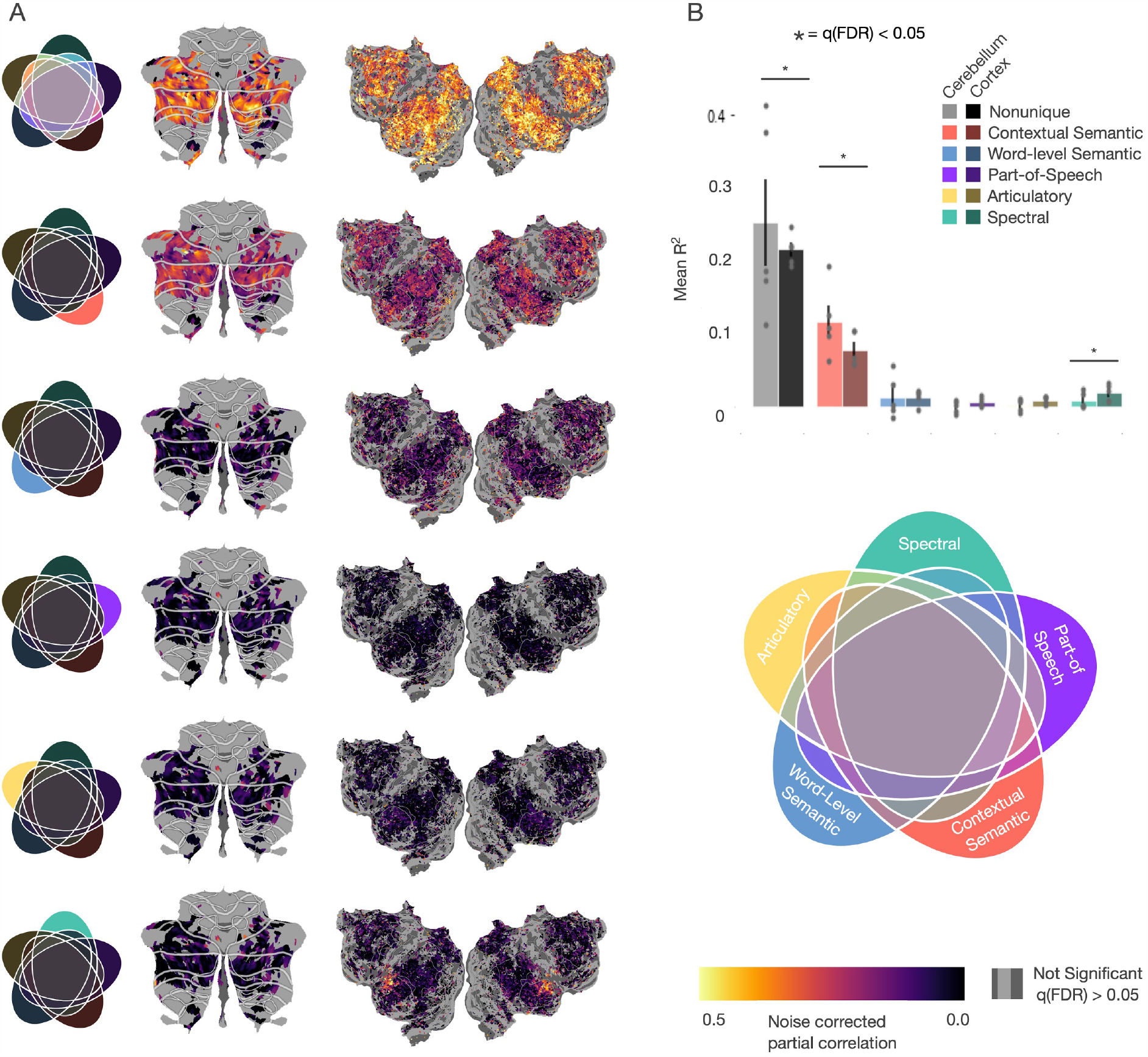
Unique variance explained by each feature space adjusted for differences in signal-to-noise in cerebellum and cortex. To determine the unique variance explained by each feature space, six new encoding models were fit: a union encoding model containing a concatenation of all feature spaces, and five encoding models each containing a concatenation of four of the five feature spaces. The unique contribution of each feature space was then determined by subtracting the variance explained by the four-way concatenation model without that feature space from the union model. This shows how much variance can be explained by each feature space above and beyond the other four. Additionally, the amount of non-unique variance—i.e., that which can be explained by more than one feature space—was determined by subtracting the 5 unique variances from the union. To account for differences in signal-to-noise between cerebellum and cortex, the correlations were corrected using standard noise-ceiling correction techniques. (**A**) The voxelwise partial correlation 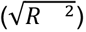 for each feature space for subject UT-S-02 was projected onto the cortical and cerebellar surfaces (Similar maps for additional subjects can be found in **Supplemental figure 5)**. Only voxels that were significantly predicted (one-sided permutation test, q(FDR)<0.05) by the 5-way union model are displayed. (**B**) Mean correlations for significant voxels in the cerebellum and cortex across all subjects. The non-unique partition explains the most variance in both cortex (darker) and cerebellum (lighter), although the variance explained by the non-unique partition is significantly (two-sided permutation test, q(FDR)<0.05) larger in cerebellum. The modality-specific spectral feature space explains significantly less variance in the cerebellum as compared to the cortex. Additionally, the modality-specific and language-specific feature spaces explain a negligible amount of variance in the cerebellum and the context-level semantic space explains the most variance among the unique partitions in cerebellum.This further supports the hypothesis that the cerebellum is largely representing language at a high, conceptual level.

**Supplemental Figure 5.**
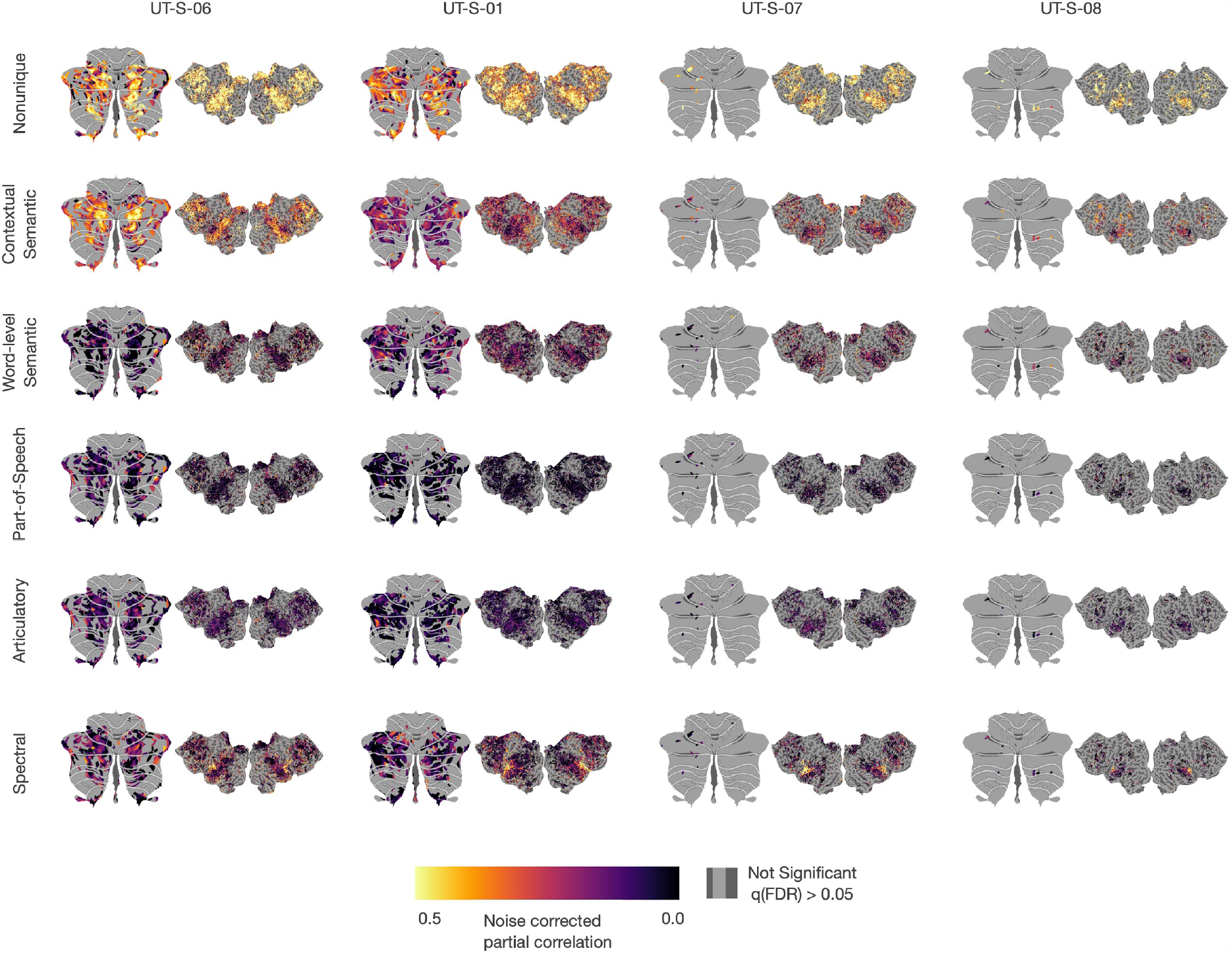
Unique variance explained for each feature space for additional subjects adjusted for differences in signal-to-noise in Cerebellum and Cortex. To determine the unique variance explained by each feature space, a union encoding model was fit with a concatenation of all feature spaces in addition to five other encoding models - each a concatenation of four of the feature spaces. The unique contribution of a model can be determined by the subtraction of the four way concatenation model without that feature space from the five way union model. Additionally, the amount of overlap between the feature spaces can be characterized by the non unique partition. The unique variance explained by each feature space in each additional subject were projected onto the cortical and cerebellar surfaces. Only significant (one-sided permutation test, q(FDR)<0.05) voxels from the 5-way joint model are displayed.

**Supplemental Figure 6.**
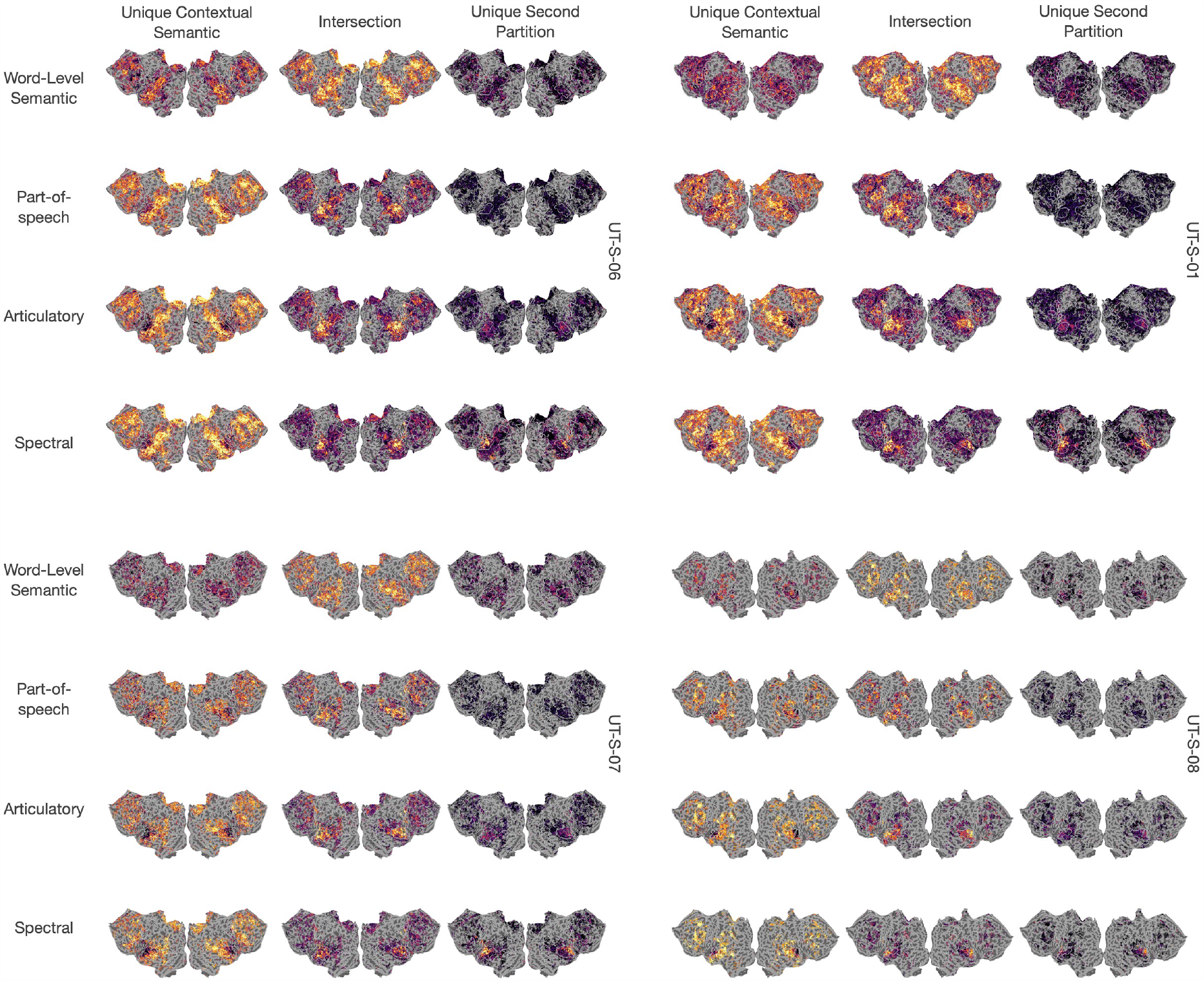
Shared explained variance of the context-level semantic feature space with each of the other feature spaces for each additional subject. To quantify the amount of overlap between the context-level semantic feature space with each of the four other feature spaces, three models for each pair of feature spaces were fit which included the concatenated feature space and each feature space individually. For each pair of models, the variance explained by each partition in each voxel was projected onto the corresponding cortical flatmaps (**Supplemental Figure 9**, noise-ceiling corrected version to account for differences in signal-to-noise across the brain) Only voxels that were significantly predicted (one-sided permutation test, q(FDR)<0.05) by each union model are shown.There is substantially lower variance explained by the intersection between the context-level semantic model and the language and modality-specific feature spaces in the cerebellum than in cortex. The results are largely consistent across subjects.

**Supplemental Figure 7.**
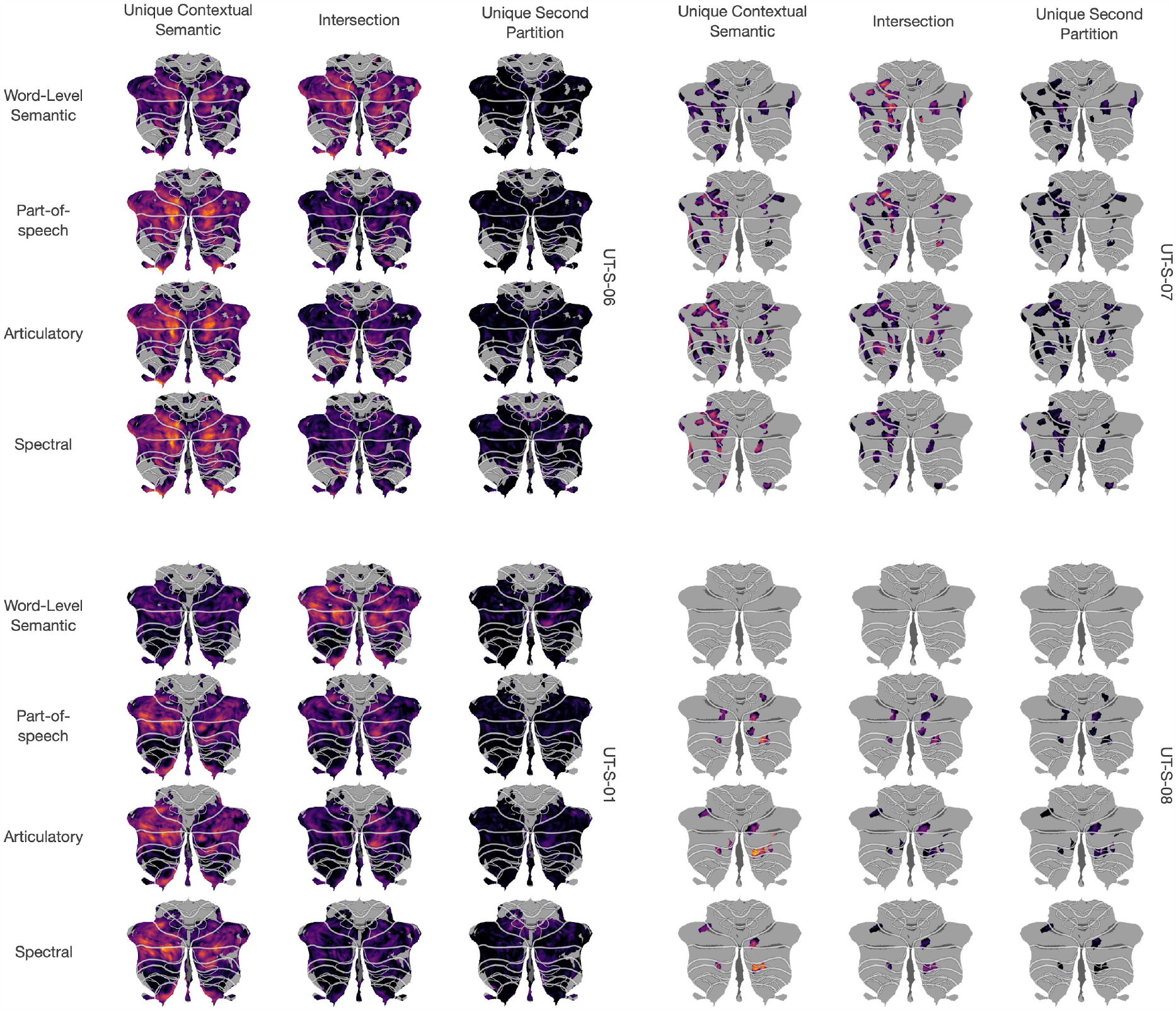
Shared explained variance of the context-level semantic feature space with each of the other feature spaces. To quantify the amount of overlap between the context-level semantic feature space with each of the four other feature spaces, three models for each pair of feature spaces were fit which included the concatenated feature space and each feature space individually. For each pair of models, the variance explained by each partition in each voxel was projected onto the corresponding cerebellar flatmaps (**Supplemental Figure 8**, noise-ceiling corrected version to account for differences in signal-to-noise across the brain) Only voxels that were significantly predicted (one-sided permutation test, q(FDR),0.05) by each union model are shown.There is substantially lower variance explained by the intersection between the context-level semantic model and the language and modality-specific feature spaces in the cerebellum than in cortex. Additionally the unique contributions for these models in the cerebellum is approaching zero and is not spatially localized. This lack of spatial localization further supports that there is no hierarchy of language processing in the cerebellum and these results provide strong support for the hypothesis that the cerebellum only represents high level, conceptual features of language, rather than low level features. This pattern of results appears consistent across subjects

**Supplemental Figure 8.**
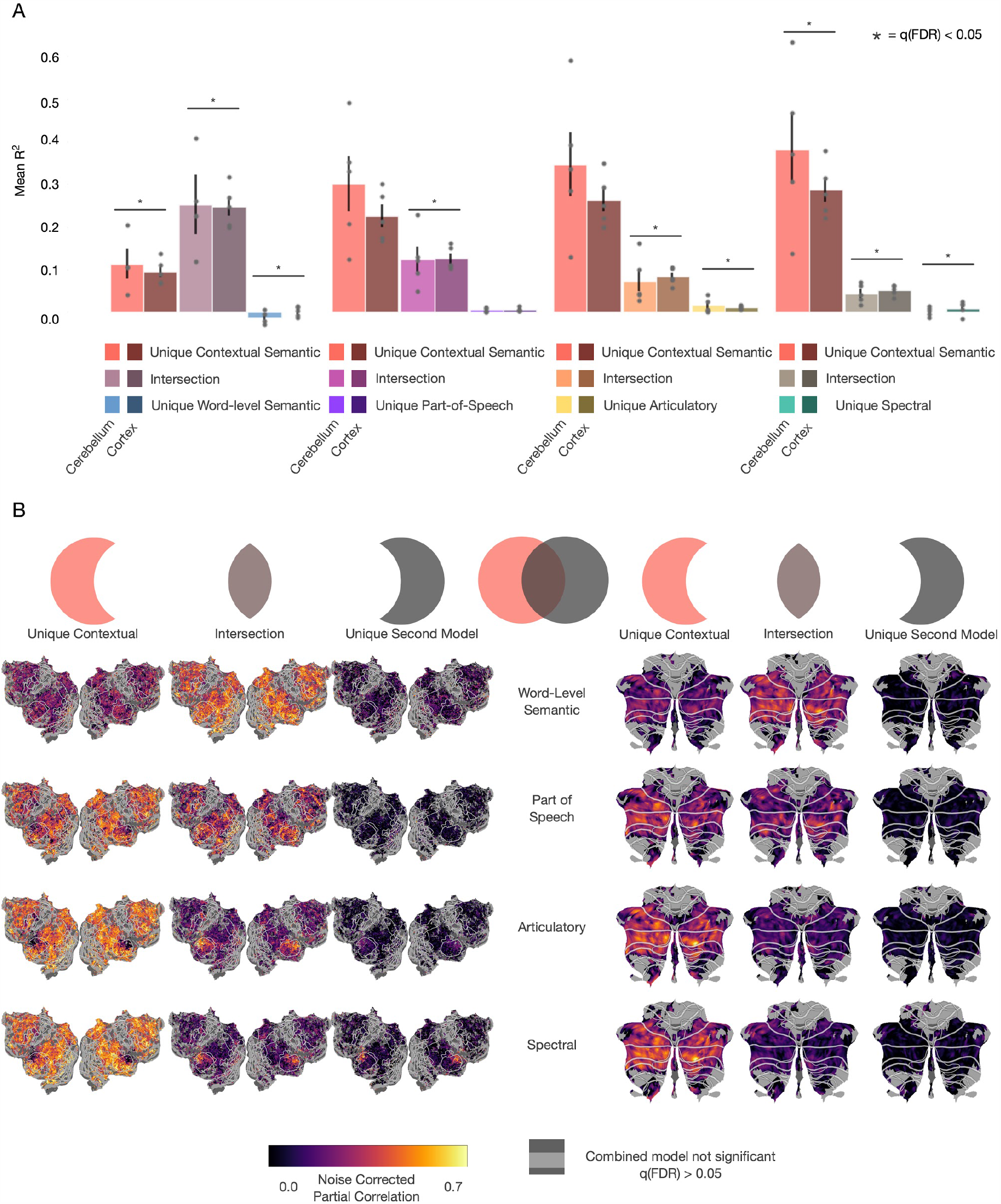
Shared explained variance of the context-level semantic feature space with each of the other feature spaces after correcting for signal-to-noise differences. To quantify the amount of overlap between the context-level semantic feature space with each of the four other feature spaces, three models for each pair of feature spaces were fit which included the concatenated feature space and each feature space individually. (**A**) For each pair models, the variance uniquely explained by the context-level feature space, uniquely explained by the second feature space, and the intersection between the two for all subjects is compared between the cerebellum and cortex. The unique context-level partition is larger in the cerebellum than in the cortex for all feature spaces (two-sided permutation test, q(FDR)<0.05). Additionally, the unique partition for each second feature space is significantly smaller in cerebellum than in cortex for every space. This shows that the lower level feature spaces predict less unique variance in the cerebellum than cortex and further supports the hypothesis that the cerebellum is not representing modality-specific or language-specific information. (**B**) For each pair of models, the variance explained by each partition in each voxel was projected onto the corresponding cortical and cerebellar flatmaps. These results are noise-ceiling corrected to account for differences in signal-to-noise ratios across the brain. Only voxels that were significantly predicted (one-sided permutation test, q(FDR)<0.05) by each union model are shown.There is substantially lower variance explained by the intersection between the context-level semantic model and the language and modality-specific feature spaces in the cerebellum than in cortex. Additionally the unique contributions for these models in the cerebellum is approaching zero and is not spatially localized. This lack of spatial localization further supports that there is no hierarchy of language processing in the cerebellum and these results provide strong support for the hypothesis that the cerebellum only represents high level, conceptual features of language, rather than low level features.

**Supplemental Figure 9.**
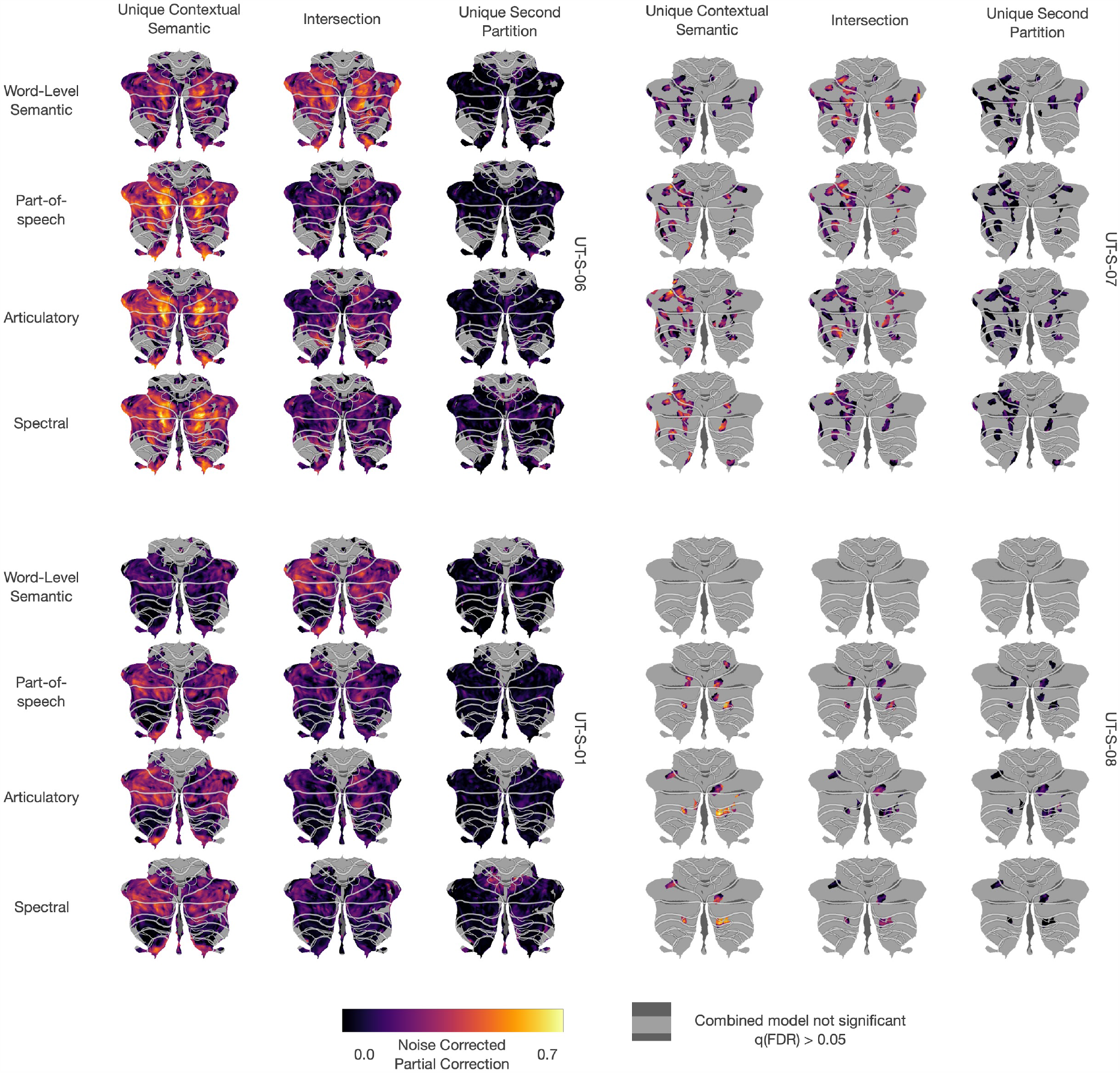
Shared explained variance of the context-level semantic feature space with each of the other feature spaces corrected for differences in signal-to-noise in cerebellum. To quantify the amount of overlap between the context-level semantic feature space with each of the four other feature spaces, three models for each pair of feature spaces were fit which included the concatenated feature space and each feature space individually. Correlations were corrected using standard noise-ceiling correction techniques to account for differences in signal-to-noise in cerebellum and cortex. For each pair of models, the variance explained by each partition in each voxel was projected onto the corresponding cerebellar flatmaps. Only voxels that were significantly predicted (one-sided permutation test, q(FDR)<0.05) by each union model are shown.There is substantially lower variance explained by the intersection between the context-level semantic model and the language and modality-specific feature spaces in the cerebellum than in cortex. Additionally the unique contributions for these models in the cerebellum is approaching zero and is not spatially localized. This lack of spatial localization further supports that there is no hierarchy of language processing in the cerebellum and these results provide strong support for the hypothesis that the cerebellum only represents high level, conceptual features of language, rather than low level features.

**Supplemental Figure 10.**
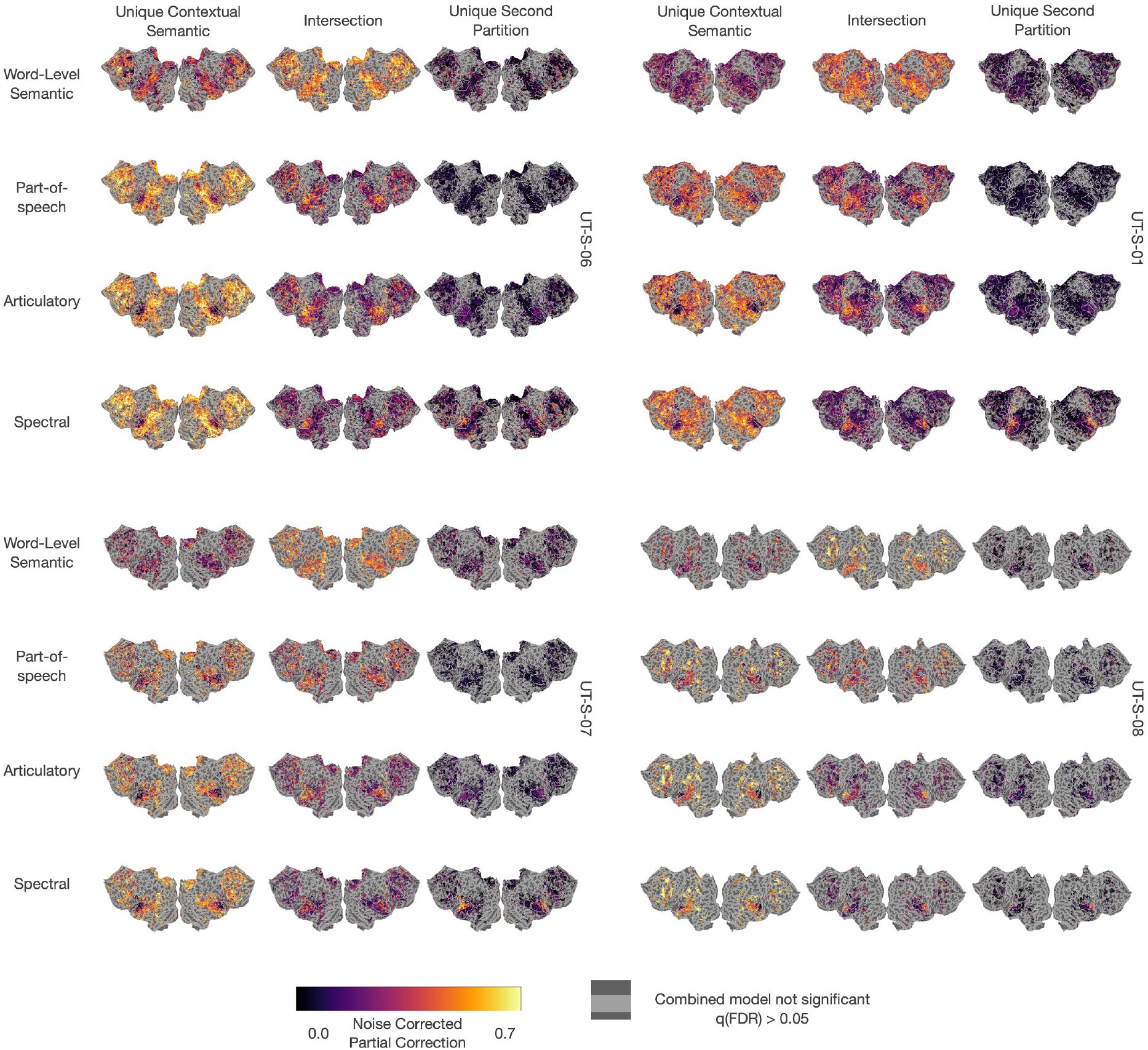
Shared explained variance of the context-level semantic feature space with each of the other feature spaces corrected for differences in signal-to-noise in cortex. To quantify the amount of overlap between the context-level semantic feature space with each of the four other feature spaces, three models for each pair of feature spaces were fit which included the concatenated feature space and each feature space individually. For each pair of models, the variance explained by each partition in each voxel was projected onto the corresponding cortical flatmaps. Correlations were corrected using standard noise-ceiling correction techniques to account for variance in the signal-to-noise ration. Only voxels that were significantly predicted (one-sided permutation test, q(FDR)<0.05) by each union model are shown.

**Supplemental Figure 11.**
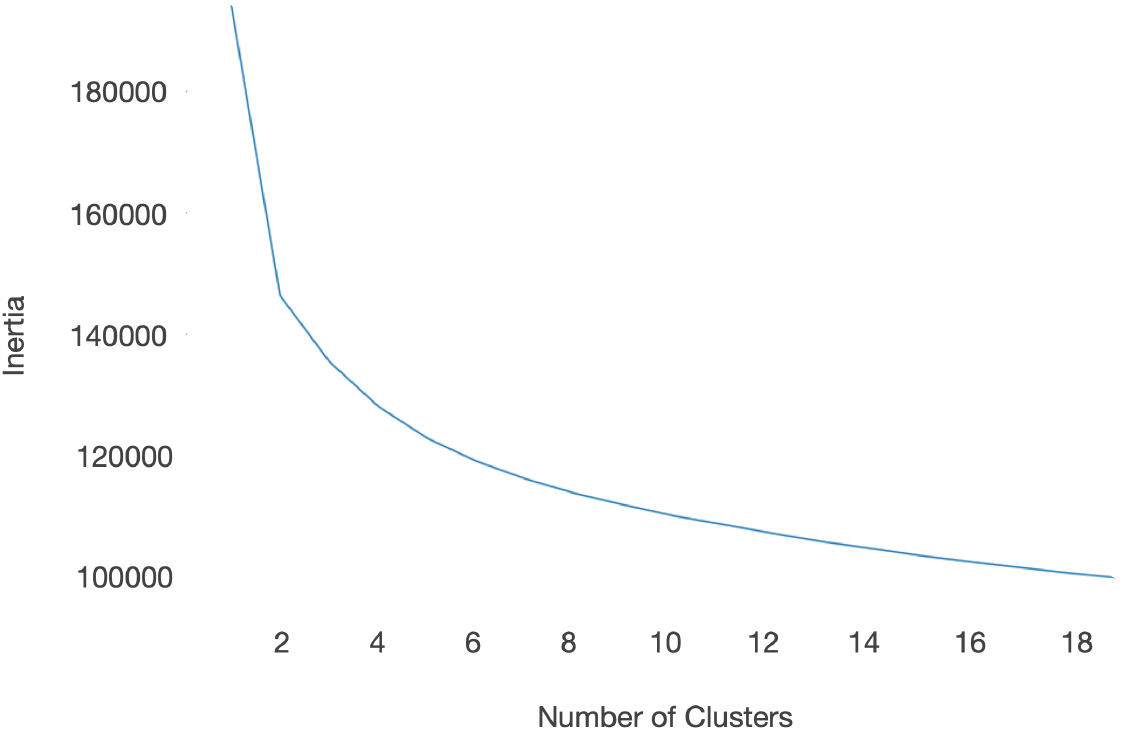
Inertia of spherical K-means clustering with increasing number of clusters. To determine the ideal number of clusters to use when clustering the model weights of word-level semantic models, we calculated the inertia at each cluster amount from 1 to 20. There is a clear “elbow point” where the slope of the inertia changes from exponential to linear at 5 clusters, thus that is what we chose to use. However, we did not find any differences in the results using a different number of clusters.

**Supplemental Figure 12.**
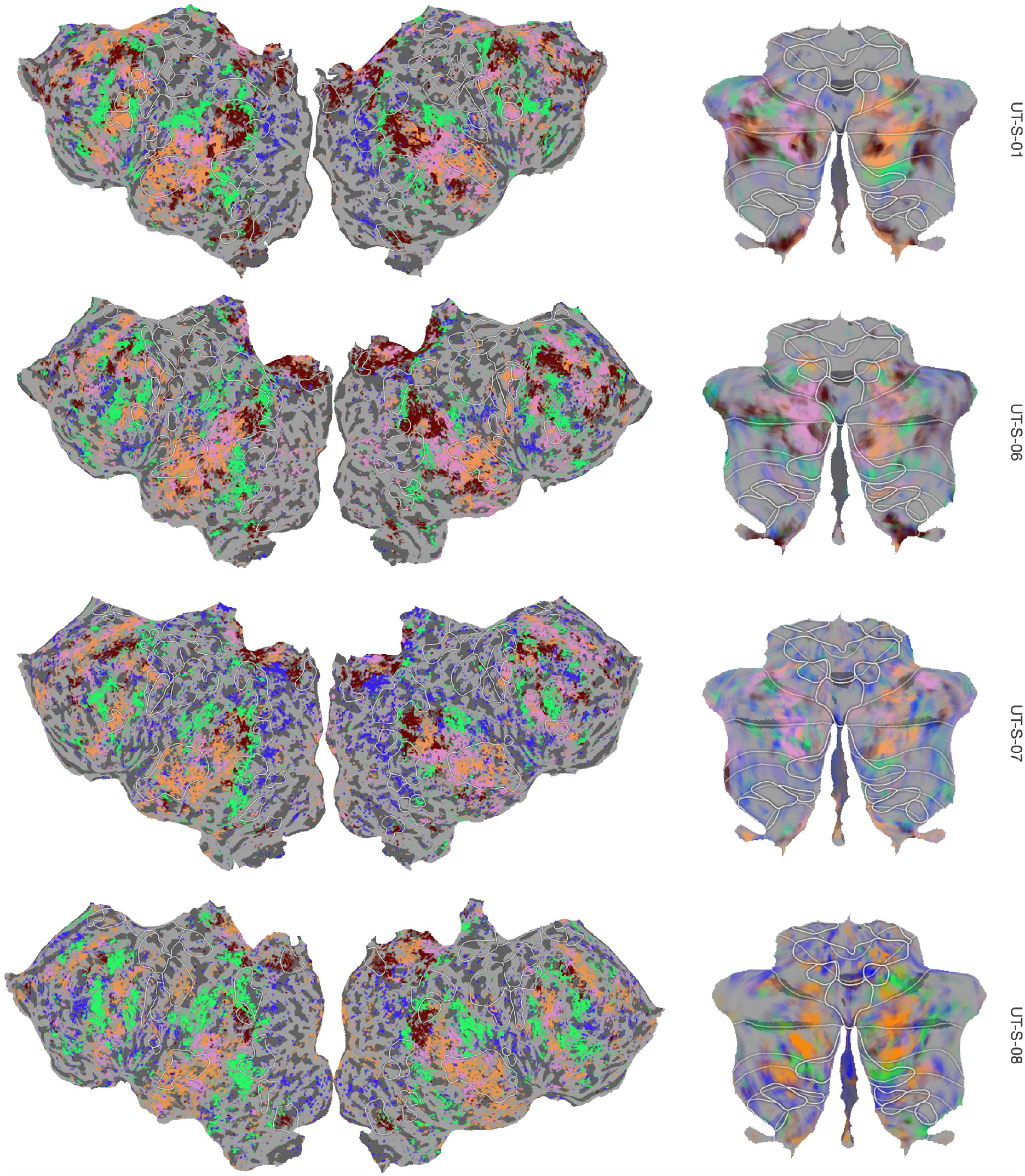
Semantic Clustering of Model Weights. To check for differences in semantic representations between the cerebellum and cortex, word-level encoding model weights from both cerebellum and cortex in all subjects were concatenated, including only the top 20% best-predicted voxels. This matrix was then clustered using spherical k-means into 5 clusters, which fell at the inflection point in the inertia graph (**Supplemental Figure 11**). For visualization, the centroid for each cluster was transformed into the same RGB space used in **Figure 5**, and each voxel in that cluster was assigned that color. The cluster distribution for each subject across the cerebellum and cortex are shown.

## References

Akbik, A., Bergmann, T., Blythe, D., Rasul, K., Schweter, S., & Vollgraf, R. (2019). FLAIR: An Easy-to-Use Framework for State-of-the-Art NLP. Proceedings of the 2019 Conference of the North American Chapter of the Association for Computational Linguistics (Demonstrations), 54–59.

Allen, G., Buxton, R. B., Wong, E. C., & Courchesne, E. (1997). Attentional activation of the cerebellum independent of motor involvement. Science, 275(5308), 1940–1943.

Berger, A. L., Della Pietra, V.J., & Della Pietra, S.A. (1996). A Maximum Entropy Approach to Natural Language Processing. Computational Linguistics, 22(1), 39–68.

Binder, J. R., Desai, R. H., Graves, W. W., & Conant, L. L. (2009). Where Is the Semantic System? A Critical Review and Meta-Analysis of 120 Functional Neuroimaging Studies. Cerebral Cortex, 19(12), 2767–2796.

Binder, J. R., Frost, J. A., Hammeke, T. A., Cox, R. W., Rao, S. M., & Prieto, T. (1997). Human brain language areas identified by functional magnetic resonance imaging. The Journal of Neuroscience: The Official Journal of the Society for Neuroscience, 17(1), 353–362.

Booth, J. R., Wood, L., Lu, D., Houk, J. C., & Bitan, T. (2007). The role of the basal ganglia and cerebellum in language processing. Brain Research, 1133, 136–144.

Brissenden, J. A., Tobyne, S. M., Osher, D. E., Levin, E. J., Halko, M. A., & Somers, D. C. (2018). Topographic Cortico-cerebellar Networks Revealed by Visual Attention and Working Memory. Current Biology: CB, 28(21), 3364–3372.e5.

Buckner, R. L., Krienen, F. M., Castellanos, A., Diaz, J. C., & Yeo, B. T. T. (2011). The organization of the human cerebellum estimated by intrinsic functional connectivity. Journal of Neurophysiology, 106(5), 2322–2345.

Caesar, K., Gold, L., & Lauritzen, M. (2003). jContext sensitivity of activity-dependent increases in cerebral blood flow. Proceedings of the Natural Academy of Science, 100(7), 4239–4244.

Callan, D. E., Tsytsarev, V., Hanakawa, T., Callan, A. M., Katsuhara, M., Fukuyama, H., & Turner, R. (2006). Song and speech: brain regions involved with perception and covert production. NeuroImage, 31(3), 1327–1342.

Chang, S.-E., Horwitz, B., Ostuni, J., Reynolds, R., & Ludlow, C. L. (2011). Evidence of left inferior frontal-premotor structural and functional connectivity deficits in adults who stutter. Cerebral Cortex, 21(11), 2507–2518.

Cheung, C., Hamiton, L. S., Johnson, K., & Chang, E. F. (2016). The auditory representation of speech sounds in human motor cortex. eLife, 5. https://doi.org/10.7554/eLife.12577

Cook, M., Murdoch, B., Cahill, L., & Whelan, B. (2004). Higher-level language deficits resulting from left primary cerebellar lesions. Aphasiology, 18(9), 771–784.

Dale, A. M., Fischl, B., & Sereno, M. I. (1999). Cortical surface-based analysis. I. Segmentation and surface reconstruction. NeuroImage, 9(2), 179–194.

de Heer, W. A., Huth, A. G., Griffiths, T. L., Gallant, J. L., & Theunissen, F. E. (2017). The Hierarchical Cortical Organization of Human Speech Processing. The Journal of Neuroscience: The Official Journal of the Society for Neuroscience, 37(27), 6539–6557.

Deniz, F., Nunez-Elizalde, A. O., Huth, A. G., & Gallant, J. L. (2019). The Representation of Semantic Information Across Human Cerebral Cortex During Listening Versus Reading Is Invariant to Stimulus Modality. The Journal of Neuroscience: The Official Journal of the Society for Neuroscience, 39(39), 7722–7736.

Diedrichsen, J. (2006). A spatially unbiased atlas template of the human cerebellum. https://doi.org/10.1016/j.neuroimage.2006.05.056

Diedrichsen, J., King, M., Hernandez-Castillo, C., Sereno, M., & Ivry, R. B. (2019). Universal Transform or Multiple Functionality? Understanding the Contribution of the Human Cerebellum across Task Domains. Neuron, 102(5), 918–928.

Downing, P. E., Jiang, Y., Shuman, M., & Kanwisher, N. (2001). A cortical area selective for visual processing of the human body. Science, 293(5539), 2470–2473.

Dronkers, N. F., Wilkins, D. P., Van Valin, R. D., Jr, Redfern, B. B., & Jaeger, J. J. (2004). Lesion analysis of the brain areas involved in language comprehension. Cognition, 92(1-2), 145–177.

Epstein, R., & Kanwisher, N. (1998). A cortical representation of the local visual environment. Nature, 392(6676), 598–601.

Fedorenko, E., Behr, M. K., & Kanwisher, N. (2011). Functional specificity for high-level linguistic processing in the human brain. Proceedings of the National Academy of Sciences of the United States of America, 108(39), 16428–16433.

Fedorenko, E., Duncan, J., & Kanwisher, N. (2013). Broad domain generality in focal regions of frontal and parietal cortex. Proceedings of the National Academy of Sciences of the United States of America, 110(41), 16616–16621.

Fiez, J. A., Petersen, S. E., Cheney, M. K., & Raichle, M. E. (1992). Impaired non-motor learning and error detection associated with cerebellar damage. A single case study. Brain: A Journal of Neurology, 115 Pt 1, 155–178.

Firth, & R J. (1957). A synopsis of linguistic theory, 1930-1955. Studies in Linguistic Analysis. https://ci.nii.ac.jp/naid/10020680394/

Frank, B., Schoch, B., Hein-Kropp, C., Hövel, M., Gizewski, E. R., Karnath, H.-O., & Timmann, D. (2008). Aphasia, neglect and extinction are no prominent clinical signs in children and adolescents with acute surgical cerebellar lesions. Experimental Brain Research. Experimentelle Hirnforschung. Experimentation Cerebrale, 184(4), 511–519.

Gao, J. S., Huth, A. G., Lescroart, M. D., & Gallant, J. L. (2015). Pycortex: an interactive surface visualizer for fMRI. Frontiers in Neuroinformatics, 9, 23.

Herculano-Houzel, S. (2010). Coordinated scaling of cortical and cerebellar numbers of neurons. Frontiers in Neuroanatomy, 4(12). https://doi.org/10.3389/fnana.2010.00012

Hickok, G., & Poeppel, D. (2007). The cortical organization of speech processing. Nature Reviews. Neuroscience, 8(5), 393–402.

Huth, A. G., de Heer, W. A., Griffiths, T. L., Theunissen, F. E., & Gallant, J. L. (2016). Natural speech reveals the semantic maps that tile human cerebral cortex. Nature, 532(7600), 453–458.

Huth, A. G., Nishimoto, S., Vu, A. T., & Gallant, J. L. (2012). A continuous semantic space describes the representation of thousands of object and action categories across the human brain. Neuron, 76(6), 1210–1224.

Jain, S., & Huth, A. G. (2018). Incorporating context into language encoding models for fMRI (Vol. 2018-Decem, pp. 6628–6637). http://papers.nips.cc/paper/7897-incorporating-context-into-language-encoding-models-for-fmri.pdf

Jiahong Yuan, M. L. (2008). Speaker identification on the SCOTUS corpus. In Proceedings of Acoustics 2008. http://citeseerx.ist.psu.edu/viewdoc/summary?doi=10.1.1.227.6546

Jürgens, U. (2002). Neural pathways underlying vocal control. Neuroscience and Biobehavioral Reviews, 26(2), 235–258.

Justus, T. (2004). The cerebellum and English grammatical morphology: evidence from production, comprehension, and grammaticality judgments. Journal of Cognitive Neuroscience, 16(7), 1115–1130.

Kanwisher, N., McDermott, J., & Chun, M. M. (1997). The fusiform face area: a module in human extrastriate cortex specialized for face perception. The Journal of Neuroscience: The Official Journal of the Society for Neuroscience, 17(11), 4302–4311.

Kawato, M., & Gomi, H. (1992). A computational model of four regions of the cerebellum based on feedback-error learning. Biological Cybernetics, 68(2), 95–103.

Kelley, E., Paul, J. J., Fein, D., & Naigles, L. R. (2006). Residual Language Deficits in Optimal Outcome Children with a History of Autism. Journal of Autism and Developmental Disorders, 36(6), 807–828.

King, M., Hernandez-Castillo, C. R., Poldrack, R. A., Ivry, R. B., & Diedrichsen, J. (2019). Functional boundaries in the human cerebellum revealed by a multi-domain task battery. Nature Neuroscience, 22(8), 1371–1378.

King, M., Hernandez-castillo, C. R., Poldrack, R., & Ivry, R. B. (2018). A Multi-Domain Task Battery Reveals Functional Boundaries in the Human Cerebellum. Neuron, 1–41.

Leiner, H. C., Leiner, A. L., & Dow, R. S. (1986). Does the cerebellum contribute to mental skills? Behavioral Neuroscience, 100(4), 443–454.

Lescroart, M. D., Stansbury, D. E., & Gallant, J. L. (2015). Fourier power, subjective distance, and object categories all provide plausible models of BOLD responses in scene-selective visual areas. Frontiers in Computational Neuroscience, 9, 135.

Levelt, W. J. M. (1993). Speaking: From Intention to Articulation. MIT Press.

Lin, Y., Tan, Y. C., & Frank, R. (2019). Open Sesame: Getting Inside BERT’s Linguistic Knowledge. In arXiv [cs.CL]. arXiv. http://arxiv.org/abs/1906.01698

Liu, L., Loannides^, A. A., & Strait, M. (1999). Single Trial Analysis of Neurophysiological Correlates of the Recognition of Complex Objects and Facial Expressions of Emotion (Vol. 11).

Manto, M., Bower, J. M., Conforto, A. B., Delgado-García, J. M. Suzete &., Farias Da Guarda, N., Gerwig, M., Habas, C., Hagura, N., Ivry, R. B., Mariën, P., Molinari, M., Naito, E., Nowak, D. A., Oulad, N., Taib, B., Pelisson, D., Tesche, C. D., Tilikete, C., … Naito, E. (2012). Consensus Paper: Roles of the Cerebellum in Motor Control-The Diversity of Ideas on Cerebellar Involvement in Movement. Cerebellum, 11, 457–487.

Marek, S., Siegel, J. S., Gordon, E. M., Raut, R. V., Gratton, C., Newbold, D. J., Ortega, M., Laumann, T. O., Adeyemo, B., Miller, D. B., Zheng, A., Lopez, K. C., Berg, J. J., Coalson, R. S., Nguyen, A. L., Dierker, D., Van, A. N., Hoyt, C. R., McDermott, K. B., … Dosenbach, N. U. F. (2018). Spatial and Temporal Organization of the Individual Human Cerebellum. Neuron, 100(4), 977–993.e7.

Mariën, P., & Manto, M. (2018). Cerebellum as a Master-Piece for Linguistic Predictability. Cerebellum, 17(2), 101–103.

Mathiesen, C., Caesar, K., & Lauritzen, M. (2000). Temporal coupling between neuronal activity and blood flow in rat cerebellar cortex as indicated by field potential analysis. The Journal of Physiology, 523(1), 235–246.

Murdoch, B. E. (2010). The cerebellum and language: Historical perspective and review. Cortex; a Journal Devoted to the Study of the Nervous System and Behavior, 46(7), 858–868.

Noppeney, U., & Price, C. J. (2004). Retrieval of abstract semantics. NeuroImage, 22(1), 164– 170.

P Boersma, D. W. (2014). Praat: doing phonetics by computer.

Petacchi, A., Laird, A. R., Fox, P. T., & Bower, J. M. (2005). Cerebellum and auditory function: an ALE meta-analysis of functional neuroimaging studies. Human Brain Mapping, 25(1), 118–128.

Petersen, S. E., Fox, P. T., Posner, M. I., Minton, M., & Raichle, M. E. (1988). Positron emission tomographic studies of the cortical anatomy of single-word processing. Nature, 331. https://www.nature.com/articles/331585a0.pdf

Poeppel, D., Emmorey, K., Hickok, G., & Pylkkänen, L. (2012). Towards a new neurobiology of language. The Journal of Neuroscience: The Official Journal of the Society for Neuroscience, 32(41), 14125–14131.

Radford, A., Narasimhan, K., Salimans, T., & Sutskever, I. (n.d.). Improving Language Understanding by Generative Pre-Training. http://s3-us-west-2.amazonaws.com/openai-assets/research-covers/language-unsupervised/language_understanding_paper.pdf

Radford, A., Narasimhan, K., Salimans, T., & Sutskever, I. (2018). Improving language understanding by generative pre-training. OpenAI. https://www.cs.ubc.ca/~amuham01/LING530/papers/radford2018improving.pdf

Radford, A., Wu, J., Child, R., Luan, D., Amodei, D., & Sutskever, I. (2019). Language models are unsupervised multitask learners. OpenAI Blog, 1(8), 9.

Schmahmann, J. D., & Sherman, J. C. (1998). The cerebellar cognitive affective syndrome. Brain: A Journal of Neurology, 121(4), 561–579.

Schoppe, O., Harper, N. S., Willmore, B. D. B., King, A. J., & Schnupp, J. W. H. (2016). Measuring the Performance of Neural Models. Frontiers in Computational Neuroscience, 10, 10.

Silveri, M. C., Leggio, M. G., & Molinari, M. (1994). The cerebellum contributes to linguistic production: a case of agrammatic speech following a right cerebellar lesion. Neurology, 44(11), 2047–2050.

Silveri, M. C., & Misciagna, S. (2000). Language, memory, and the cerebellum. Journal of Neurolinguistics, 13(2), 129–143.

Snider, R. S., & Stowell, A. (1944). RECEIVING AREAS OF THE TACTILE, AUDITORY, AND VISUAL SYSTEMS IN THE CEREBELLUM. Journal of Neurophysiology, 7(6), 331–357.

Stoodley, C. J., & Schmahmann, J. D. (2009). The cerebellum and language: Evidence from patients with cerebellar degeneration. Brain and Language, 110(3), 149–153.

Tenney, I., Xia, P., Chen, B., Wang, A., Poliak, A., Thomas McCoy, R., Kim, N., Van Durme, B., Bowman, S. R., Das, D., & Pavlick, E. (2019). What do you learn from context? Probing for sentence structure in contextualized word representations. In arXiv [cs.CL]. arXiv. http://arxiv.org/abs/1905.06316

Toneva, M., & Wehbe, L. (2019). Interpreting and improving natural-language processing (in machines) with natural language-processing (in the brain). Advances in Neural Information Processing Systems. http://papers.nips.cc/paper/9633-interpreting-and-improving-natural-language-processing-in-machines-with-natural-language-processing-in-the-brain

Vaishnavi, S. N., Vlassenko, A. G., Rundle, M. M., Snyder, A. Z., Mintun, M. A., & Raichle, M. E. (2010). Regional aerobic glycolysis in the human brain. Proceedings of the National Academy of Science, 107(41), 17757–17762.

Woolrich, M. W., Jbabdi, S., Patenaude, B., Chappell, M., Makni, S., Behrens, T., Beckmann, C., Jenkinson, M., & Smith, S. M. (2009). Bayesian analysis of neuroimaging data in FSL. NeuroImage, 45(1 Suppl), S173–S186.

